# NTMC2T5 links lipid homeostasis to plastid differentiation

**DOI:** 10.64898/2026.08.26.747221

**Authors:** Carolina Huercano, Oliver Cuevas, Paula Velasco-Palomo, Miriam Moya-Barrientos, Francisco Percio, Joaquin J. Salas-Liñan, Victoria Sanchez-Vera, Noemi Ruiz-Lopez

## Abstract

Chloroplast biogenesis requires extensive lipid remodeling to establish the internal membrane systems of developing plastids, yet how lipid homeostasis is coordinated during this process remains incompletely understood. Here, we identify a previously unrecognized, Archaeplastida-conserved family of SMP-domain proteins and characterize its role in early plastid development. NTMC2T5 proteins contain an N-terminal chloroplast-targeting membrane region, an SMP domain, and a C2 domain, and localize in punctate patterns at the chloroplast envelope, enriched at regions associated with the endoplasmic reticulum (ER). Loss of NTMC2T5 in *Nicotiana benthamiana* causes severe defects in chloroplast development during seedling establishment and de-etiolation, whereas chloroplast maintenance in mature leaves is largely unaffected. Ultrastructural analyses revealed that mutant plastids fail to establish normal prolamellar bodies and organized thylakoid membranes, although plastid number and size were largely unaffected. Lipidomic analyses further revealed that NTMC2T5 loss causes a strong reduction in the plastid galactolipids monogalactosyldiacylglycerol and digalactosyldiacylglycerol, accompanied by accumulation of extraplastidial phospholipids and altered fatty-acid composition during de-etiolation. Together, these findings identify NTMC2T5 as a previously unrecognized determinant of lipid homeostasis during plastid differentiation and establish a link between a plant-specific SMP-domain protein family and chloroplast membrane biogenesis. We propose that NTMC2T5 contributes to ER–plastid lipid exchange and/or organization of ER–plastid membrane interfaces during early chloroplast development.

**HIGHLIGHTS:**

- NTMC2T5 defines a previously unrecognized SMP-domain protein family conserved across Archaeplastida.
- NTMC2T5 localizes to punctate chloroplast-envelope regions associated with the endoplasmic reticulum.
- NTMC2T5 is required for plastid differentiation during early seedling development.
- Loss of NTMC2T5 disrupts internal plastid membrane formation and lipid remodeling.
- NTMC2T5 links lipid homeostasis to plastid differentiation.

## INTRODUCTION

Chloroplast biogenesis is a fundamental developmental process that enables young seedlings to transition from heterotrophic growth sustained by seed reserves to autotrophic growth driven by photosynthesis (Albrecht *et al*., 2006; Liebers *et al*., 2022). During this transition, undifferentiated proplastids in seeds differentiate into functional chloroplasts in cotyledons (Pogson and Albrecht, 2011). The establishment of photosynthetic competence requires coordinated expression of nuclear and plastid genes, assembly of photosynthetic protein complexes, and extensive membrane biogenesis (Kakizaki *et al*., 2009; Pipitone *et al*., 2021). These processes occur within a relatively narrow developmental window and culminate in the establishment of the photosynthetic machinery.

A major component of this transition is the expansion and reorganization of the chloroplast membrane system. When germination occurs in the light, proplastids can differentiate directly into chloroplasts, whereas in darkness they develop into etioplasts containing characteristic prolamellar bodies and prothylakoid membranes that are subsequently reorganized upon illumination to generate mature thylakoids (Pogson and Albrecht, 2011). In both developmental trajectories, chloroplast maturation requires the production of large amounts of membrane material. This is accompanied by a marked accumulation of the major chloroplast galactolipids monogalactosyldiacylglycerol (MGDG) and digalactosyldiacylglycerol (DGDG) as the thylakoid system is established and expanded (Fujii *et al*., 2014; Pipitone *et al*., 2021).

Although chloroplasts possess the enzymatic machinery required for *de novo* fatty acid synthesis, their lipid metabolism is closely integrated with that of the endoplasmic reticulum (ER) (Browse *et al*., 1986; Ohlrogge and Browse, 1995). Genetic and biochemical studies have established that ER-derived lipid precursors make a substantial contribution to chloroplast membrane biogenesis, particularly during early plastid differentiation (Negi *et al*., 2018; Obata *et al*., 2021; Pipitone *et al*., 2021). This metabolic interdependence raises a fundamental cell biological question: how are lipids exchanged efficiently between the ER and developing plastids?

Membrane contact sites provide a mechanism for such inter-organelle exchange. At these specialized regions, two organelles are held in close proximity, typically within tens of nanometers, without membrane fusion. This creates a confined environment that facilitates the transfer of lipids and other molecules between them (Scorrano *et al*., 2019). Ultrastructural studies have long revealed physical associations between plastids and the ER, and recent work has established ER–plastid membrane contact sites as important hubs for inter-organelle communication (Andersson *et al*., 2007; Yao *et al*., 2023; Renna *et al*., 2024; Huercano *et al*., 2025, 2026). Several molecular components involved in lipid trafficking at these interfaces have now been identified, including the Trigalactosyldiacylglycerol (TGD1–TGD5) machinery (Xu *et al*., 2003, 2005; Awai *et al*., 2006; Lu *et al*., 2007; Roston *et al*., 2012; Fan *et al*., 2015), Sec14-like proteins SFH5 and SFH7, which can mediate phosphatidic acid transfer from ER to chloroplast (Yao *et al*., 2023), and the VAP27–ORP2A complex, which defines a functional ER-chloroplast contact site and contributes to chloroplast-envelope lipid homeostasis (Renna *et al*., 2024). Perturbation of these components affects plastid lipid homeostasis and, in several cases, chloroplast development, demonstrating the physiological importance of ER– plastid lipid exchange. Nevertheless, the molecular machinery operating at these interfaces remains incompletely defined.

A particularly important class of proteins associated with lipid transport and membrane organization at contact sites is defined by the presence of the Synaptotagmin-like Mitochondrial-lipid-binding Protein (SMP) domain. SMP domains belong to the TULIP (Tubular Lipid-binding) superfamily and adopt an elongated, tubular fold enclosing a hydrophobic cavity capable of accommodating glycerolipids (Kopec *et al*., 2010; AhYoung *et al*., 2015). Structural and biochemical studies have shown that SMP domains can bind and transfer lipids between closely apposed membranes, providing a mechanism for non-vesicular lipid transport at membrane contact sites (Kornmann *et al*., 2009; Reinisch and De Camilli, 2016; Bian and De Camilli, 2019; Jeyasimman and Saheki, 2020). SMP-domain proteins are found in diverse membrane contact-site machineries and can contribute to both lipid transfer and membrane tethering or organization. In yeast, several SMP-domain proteins are components of the ER– mitochondria encounter structure (ERMES), contributing to physical and functional coupling between the ER and mitochondria (Kornmann *et al*., 2009; Wozny *et al*., 2023). In animals, Extended Synaptotagmins and PDZD8 likewise participate in lipid exchange at distinct membrane contact sites (Giordano *et al*., 2013; Hirabayashi *et al*., 2017; Guillén-Samander *et al*., 2019; Sassano *et al*., 2023). Plant SMP-domain proteins have also been implicated in lipid homeostasis. In *Arabidopsis*, the ER-resident Synaptotagmins SYT1 and SYT3 contribute to the maintenance of plasma-membrane diacylglycerol homeostasis during abiotic stress (Ruiz-Lopez *et al*., 2021).

Whether plants possess additional SMP-domain proteins specialized for lipid exchange or membrane organization at ER–plastids contact sites remains largely unexplored. Plant Synaptotagmin 1 (SYT1) represents the best-characterized plant SMP-domain protein and has established roles at ER–plasma membrane contact sites (Yamazaki *et al*., 2009; Pérez-Sancho *et al*., 2015; Ruiz-Lopez *et al*., 2021; Garcia-Hernandez *et al*., 2025), but the broader diversity and functional specialization of SMP-domain proteins remain poorly defined. A previous analysis of nucleotide sequence databases identified several plant proteins containing SMP domains (Craxton, 2007), providing an early indication of a broader repertoire of these proteins in plants. However, a comprehensive characterization of plant SMP-domain proteins is still needed to determine their diversity, subcellular localization, and potential contributions to inter-organelle lipid exchange.

Here, we systematically identify SMP-domain proteins in *Arabidopsis* and *Nicotiana benthamiana* and show that the N-terminal transmembrane C2 domain type 5 (NTMC2T5) family is conserved across Archaeplastida. We investigated the subcellular localization and molecular properties of NTMC2T5 proteins and examined their contribution to chloroplast development during seedling establishment and de-etiolation. Our findings reveal a previously unrecognized role for a plant SMP-domain protein family in plastid biogenesis and establish a link between SMP proteins, ER-associated chloroplast membrane interfaces, and the lipid homeostasis required for chloroplast membrane biogenesis.

## RESULTS

### Phylogenetic analysis identifies NTMC2T5 as a conserved SMP-domain protein family in Archaeplastida

To identify the repertoire of SMP-domain proteins in plants, we performed iterative homology searches using the sequence of the SMP region of human E-Syt1 (D135– V313) as a query. Searches were conducted using HHpred against the *Arabidopsis thaliana* proteome (TAIR10, 20 Jun 2017; database: PDB_mmCIF70_24_Dec) (Zimmermann *et al*., 2018) and PSI-BLAST against the *Nicotiana benthamiana* predicted proteome (Solgenomics v1.0.1 predicted proteins), combining the high sensitivity of profile-based detection with broader species coverage. This strategy identified 15 SMP-domain-containing proteins in *A. thaliana*—including AtSYT1-6, AtCLB1, AtNTMC2T5.1/2, AtNTMC2T6.1/2, AtTEX2A/B, At3G60950, and At3G61030—and 29 putative homologs in *N. benthamiana* (hereafter designated with the Nb prefixed).

We then performed Cluster Analysis of Sequences (CLANS) (Frickey and Lupas, 2004) to visualize relationships among SMP-domain proteins based on all-against-all pairwise sequence similarities. The dataset included all putative SMP-domain proteins from *Arabidopsis* and *N. benthamiana*, together with all identified SMP-domain proteins from humans, *Saccharomyces cerevisiae* and the SMP-domain-containing plant proteins described by Craxton (Craxton, 2007) (Fig. 1A). This analysis resolved seven major SMP-domain protein clusters in plants corresponding to SYT1–3, SYT4–5, CLB1, SYT6, NTMC2T5, NTMC2T6 and TEX2. Canonical plant synaptotagmins (SYT1-5) clustered with their metazoan orthologs, suggesting a conserved evolutionary origin across eukaryotes. In contrast, NTMC2T5, NTMC2T6, and SYT6 formed well-separated plant-specific clusters, indicating substantial diversification of SMP-domain proteins within the green lineage (Fig. 1B). TEX2 proteins constituted a highly divergent and weakly connected group (Fig. 1C). Notably, no close relationships were detected between plant SMP-domain proteins and the mammalian SMP-domain proteins TMEM24 or PDZD8, nor with yeast ERMES components (Fig. 1A-B).

**Figure 1.**
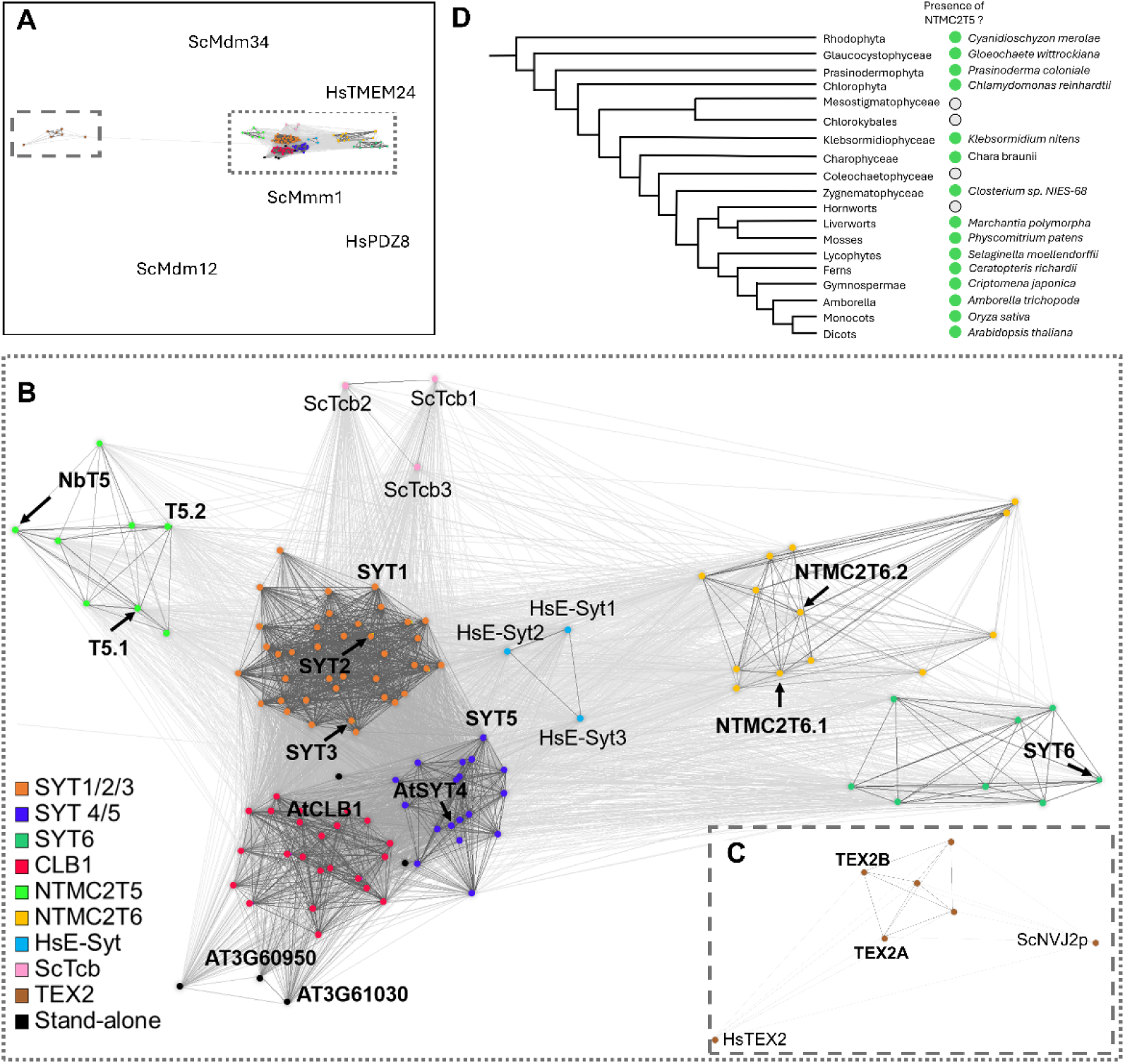
Identification of SMP-domain proteins in *Arabidopsis thaliana* and *Nicotiana benthamiana* and their evolutionary relationships across eukaryotes. **(A)** Clustering map of all SMP-domain proteins identified in this study, together with previously characterized SMP-containing proteins from *Homo sapiens* and *Saccharomyces cerevisiae* and the NTMC2 proteins identified by Craxton (2007) in a range of plant species (Supplemental File S1). Proteins were clustered based on their all-against-all pairwise sequence similarities using CLANS (Frickey and Lupas, 2004), with the BLOSUM62 matrix and an E-value cutoff of 1×10⁻⁹. Dots represent sequences, and sequences assigned to the same group share the same color. Sequences that could not be assigned to a group are labeled as standalone proteins. Lines connect sequences with detectable similarity, and line color reflects the E-value of the pairwise comparison (brighter colors indicate lower E-values). Boxed regions are magnified in (B) and (C). **(B)** Magnified view of the dotted box in (A). This region contains the SYT1–3, SYT4–5, CLB1, SYT6, NTMC2T5 and NTMC2T6 clusters, together with the *S. cerevisiae* Tcb and *H. sapiens* E-Syt proteins. HsPDZD8, HsTMEM24 and the *S. cerevisiae* ERMES components lie outside this region. **(C)** Magnified view of the dashed box in (A), showing the CLANS relationships between TEX2 proteins from *A. thaliana*, *H. sapiens* and *S. cerevisiae*. **(D)** Detection of NTMC2T5 homologs across the Archaeplastida lineage, from red algae to eudicots, mapped onto a simplified evolutionary scheme. Clades are indicated by their taxonomic name (e.g., Rhodophyta). Green circles denote clades in which at least one NTMC2T5 homolog was identified, with the corresponding species indicated next to the circle. Gray circles indicate clades in which no candidate protein was detected by PSI-BLAST searches against the corresponding taxonomic group. *Abbreviations:* At, *Arabidopsis thaliana*; AtT5.1, AtNTMC2T5.1; AtT5.2, AtNTMC2T5.2; Hs, *Homo sapiens*; Nb, *Nicotiana benthamiana*; NbT5, NbNTMC2T5; Sc, *Saccharomyces cerevisiae*.

Among the plant-specific SMP-domain groups identified, NTMC2T5 formed a well-resolved cluster (Fig. 1B). To trace the evolutionary distribution of this subfamily, we screened representative photosynthetic eukaryotes for NTMC2T5 homologs and mapped their occurrence onto a consensus phylogenetic scheme (Fig. 1D). NTMC2T5 homologs were detected throughout all major Archaeplastida lineages analyzed, including early-diverging red algae (*Cyanidioschyzon merolae*), glaucophytes (*Gloeochaete wittrockiana*), green algae (chlorophytes and charophytes) and all major land plant groups (bryophytes, lycophytes, ferns, gymnosperms, and angiosperms) (Fig. 1D; Fig. S1). This broad phylogenetic distribution is consistent with an ancient origin of the NTMC2T5 subfamily early in Archaeplastida evolution. Interestingly, the distribution of NTMC2T5 also suggests several lineage-specific secondary loss events. Homologs were not detected in Mesostigmatophyceae/Chlorokybales, Coleochaetophyceae, or hornworts, despite their presence in closely related lineages. In particular, the conservation of NTMC2T5 in both liverworts and mosses suggests that its absence in hornworts is more likely attributable to a lineage-specific loss than to an ancestral absence. Furthermore, no proteins exhibiting the characteristic NTMC2T5 domain architecture were identified in *Homo sapiens* or *Saccharomyces cerevisiae*, consistent with a specialization of this architecture within Archaeplastida. Together, these observations indicate that NTMC2T5 is an ancient and evolutionarily conserved family of SMP-domain proteins present throughout Archaeplastida.

### NTMC2T5 proteins localize to the chloroplast envelope

Plant synaptotagmins (SYTs, SYT1-SYT5) are well-established components of endoplasmic reticulum (ER)–plasma membrane contact sites. In contrast, subcellular localization predictions generated using DeepLoc 2.1 (Ødum *et al*., 2024) uniquely assigned NTMC2T5 proteins to plastids (Table S1). Predicted plastid localization was conserved across most NTMC2T5 homologs identified in our phylogenetic survey, suggesting that plastid association is an evolutionarily conserved feature of this protein family.

The Arabidopsis genome encodes two NTMC2T5 proteins, AtNTMC2T5.1 (AT1G50260) and AtNTMC2T5.2 (AT3G19830), which are currently annotated as calcium-dependent lipid-binding (CaLB domain) proteins in Araport11. In contrast, the *Nicotiana benthamiana* genome contains a single NTMC2T5 homolog (NbNTMC2T5; NIBEN101SCF08519G00002). Domain analysis predicted a conserved architecture comprising a putative N-terminal transmembrane (TM) segment, an SMP domain, and a C2 domain (Fig. 2A,B), partially resembling the organization of SYT1 (Pérez-Sancho *et al*., 2015). In addition, NTMC2T5 proteins possess a short hydrophobic region (HR) at the C terminus, consisting of 20 amino acids in Arabidopsis and Nicotiana NTMC2T5 proteins (TOPCONS predictor (Tsirigos *et al*., 2015)).

**Figure 2.**
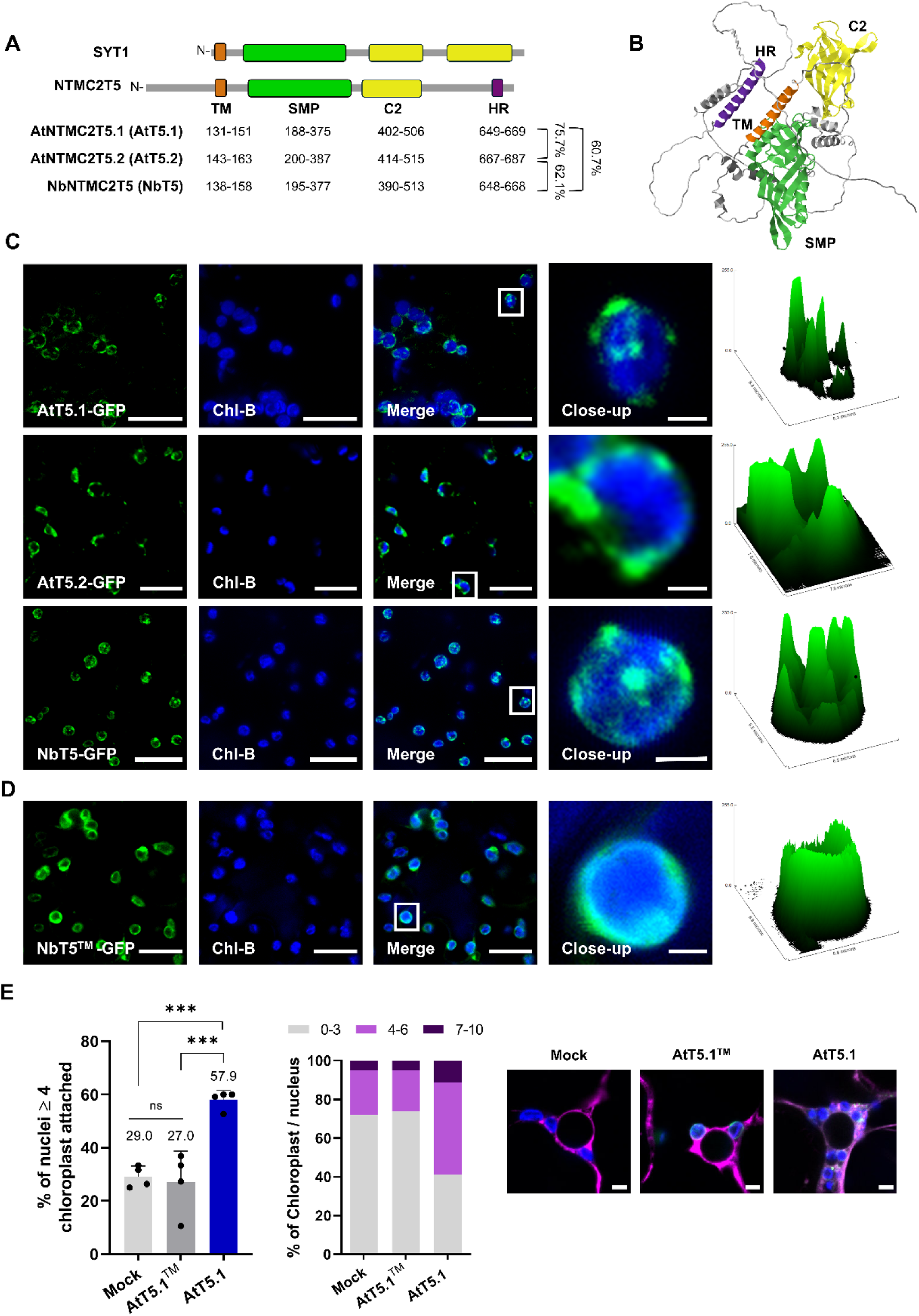
NTMC2T5 proteins localize to the chloroplast envelope. **(A)** Domain representations of SYT1 and NTMC2T5 proteins. Domains were predicted using InterPro (Blum *et al*., 2025) and TOPCONS (Tsirigos *et al*., 2015). Numbers indicate amino acid positions. Sequence similarities are shown as percentages on the right. **(B)** Three-dimensional structure of NbT5 predicted with AlphaFold3 (Abramson *et al.,* 2024). Domains are colored as in (A). **(C)** Maximum Z-projections of confocal images of *Nicotiana benthamiana* leaves transiently expressing AtT5.1-, AtT5.2- or NbT5-GFP. All three proteins localized to the chloroplast envelope and displayed a non-uniform distribution. Boxed regions are magnified in the insets. Surface plots of the fluorescence intensity of the insets (right panels) illustrate this spatial distribution. Scale bar: 20 µm. Scale bar inset: 2 µm. **(D)** Maximum Z-projections of confocal images of *N. benthamiana* leaves transiently expressing NbT5^TM^-GFP (M1-L185). NbT5^TM^-GFP uniformly localized around chloroplasts. Boxed regions are magnified in the insets. Surface plots of the fluorescence intensity of the insets (right panels) illustrate the spatial distribution of the GFP signal. Scale bar: 20 µm. Scale bar inset: 2 µm. **(E)** Quantification of chloroplast clustering around the nucleus in epidermal cells of *N. benthamiana* leaves transiently expressing ER-mCh alone (control), AtT5.1^TM^-GFP (M1-G160) or AtT5.1-GFP. ER-mCh was co-expressed in all cases to visualize nuclei. From left to right: percentage of nuclei with ≥ 4 attached chloroplasts, number of chloroplasts per nucleus, and representative single-plane confocal images containing the equatorial section of the nucleus for each condition. Merged images show chlorophyll autofluorescence (blue), GFP (green) and ER-mCh (magenta). Bars represent mean ± SD. Four plants per condition were analyzed, with approximately 20 nuclei per plant. Data were analyzed using one-way ANOVA followed by Tukey’s multiple comparison test. Asterisks indicate significant differences (ns, not significant; ***, *P* < 0.001). *Abbreviations:* AtT5.1, AtNTMC2T5.1; AtT5.2, AtNTMC2T5.2; C2, Ca²⁺ binding domain; HR, hydrophobic region; NbT5, NbNTMC2T5; SMP, Synaptotagmin-like Mitochondrial-lipid-binding Protein domain; TM, transmembrane region.

To determine the subcellular localization of NTMC2T5 proteins, C-terminal GFP fusions of Arabidopsis AtNTMC2T5.1 (AtT5.1-GFP), AtNTMC2T5.2 (AtT5.2-GFP), and *N. benthamiana* NbNTMC2T5 (NbT5-GFP) were transiently expressed in *N. benthamiana* leaves under the control of the UBIQUITIN10 promoter (UBQ10) (Grefen *et al*., 2010). Confocal imaging revealed that all three proteins localized to the chloroplast envelope and exhibited a non-uniform, punctate distribution along the chloroplast surface (Fig. 2C). Fluorescence intensity surface plots further highlighted the heterogeneous distribution of the proteins within the chloroplast envelope. To assess the role of the N-terminal putative transmembrane region in chloroplast targeting, a construct comprising the N-terminal region containing the predicted transit peptide plus the transmembrane segment (amino acids 1–160) fused to GFP (NbT5^TM^-GFP) was expressed in *N. benthamiana*. In contrast to the punctate localization of the full-length protein, NbT5^TM^-GFP produced a continuous fluorescence signal surrounding chloroplasts, consistent with localization to the chloroplast envelope (Fig. 2D). Likewise, the N-terminal transmembrane region of AtT5.1 and AtT5.2 was sufficient to direct GFP to the chloroplast periphery (AtT5.1^TM^-GFP and AtT5.2^TM^-GFP, respectively) (Fig. S2A). In contrast, deletion of the transmembrane region from AtT5.1 (AtT5.1^ΔTM^-GFP) abolished chloroplast localization and resulted in a reticulate fluorescence pattern characteristic of the ER. Co-expression with the ER marker CD3-959 (ER-mCherry) (Nelson et al., 2007) confirmed ER localization of the truncated protein (Fig. S2B,C).

Interestingly, expression of full-length AtT5.1 also affected chloroplast positioning within the cell. Whereas chloroplasts were only occasionally associated with the nucleus in mock-treated cells or in cells expressing the isolated N-terminal transmembrane region, expression of full-length AtT5.1 resulted in a pronounced accumulation of chloroplasts around the nucleus (Fig. 2E). Quantification confirmed a significant increase in the proportion of nuclei associated with four or more chloroplasts in AtT5.1-expressing cells compared with both mock and AtT5.1^TM^-expressing cells. Similarly, full-length AtT5.2 also promoted perinuclear clustering (Fig. S2D). Perinuclear chloroplast clustering is a well-established cellular response associated with chloroplast–nucleus communication and stress signaling in *N. benthamiana* (Ding *et al*., 2019). Notably, overexpression of the chloroplast outer envelope protein OMP24 has also been shown to induce perinuclear chloroplast clustering, whereas its isolated transmembrane domain, despite being sufficient for chloroplast targeting, was unable to trigger this response (Wang *et al*., 2012b; Han *et al*., 2023). Thus, the ability of full-length AtT5.1, but not its isolated transmembrane region, to promote chloroplast clustering provides additional functional support for its association with the chloroplast envelope and suggests that regions outside the transmembrane segment contribute to NTMC2T5-dependent chloroplast organization.

### NTMC2T5 is required for chloroplast biogenesis during early seedling development and de-etiolation

The localization of NTMC2T5 to the chloroplast envelope, together with the established roles of SMP-domain proteins in lipid transport and membrane homeostasis (Reinisch and De Camilli, 2016; Jeyasimman and Saheki, 2020; Huercano *et al*., 2025, 2026), prompted us to investigate whether NTMC2T5 contributes to chloroplast biogenesis. We generated CRISPR/Cas9 knockout lines targeting the first exon of *NbT5* using two independent sgRNAs (Fig. S3A,B). Two independent homozygous, transgene-free mutant lines, *nbt5 #14* and *#34*, were selected for further analysis (Fig. S3C,D). To verify that the resulting phenotypes were specifically caused by the loss of *NbT5*, we complemented *nbt5 #14* with a UBQ10::NbT5-GFP transgene.

Next, we examined seedling development under long-day conditions on half-strength MS medium. At 4–6 days after sowing, both *nbt5* mutant lines displayed pale-green to variegated cotyledons, whereas wild-type (WT) seedlings developed uniformly green cotyledons (Fig. 3A,B; Fig. S3E). Introduction of the UBQ10::NbT5-GFP transgene fully restored the WT phenotype, demonstrating that the observed developmental defects resulted specifically from disruption of *NbT5* (Fig. 3A,B; Fig. S3E). Consistent with the visual phenotype, chlorophyll accumulation was markedly reduced in *nbt5* seedlings and restored to WT levels in complemented plants (Fig. 3C; Fig. S4A). Because chlorotic and variegated cotyledon phenotypes are frequently associated with defects in chloroplast biogenesis (Miura *et al*., 2007; Zhang *et al*., 2016; Xu *et al*., 2022; Guo *et al*., 2024), we estimated chloroplast abundance in cotyledon epidermal cells using chlorophyll autofluorescence. Both mutant lines exhibited significantly reduced chlorophyll autofluorescence-positive structures relative to WT and complemented seedlings (Fig. 3D). This reduction was not rescued by exogenous sucrose supplementation, indicating that the phenotype cannot be attributed solely to impaired photosynthetic carbon assimilation (Fig. S4B). Remarkably, the requirement for *NbT5* appeared largely restricted to early developmental stages. Root growth was unaffected in 7-day-old seedlings (Fig. S4C), and as plants matured, *nbt5* mutants became phenotypically indistinguishable from WT plants (Fig. S4D). Although density of chlorophyll autofluorescence-positive structures remained lower in cotyledons at 12 and 18 days after sowing (Fig. 3E,F), no differences were detected in true leaves of age-matched plants (Fig. 3F). Likewise, adult and senescing leaves exhibited chlorophyll contents comparable to those of WT plants, as determined by SPAD measurements (Fig. S4E,F).

**Figure 3.**
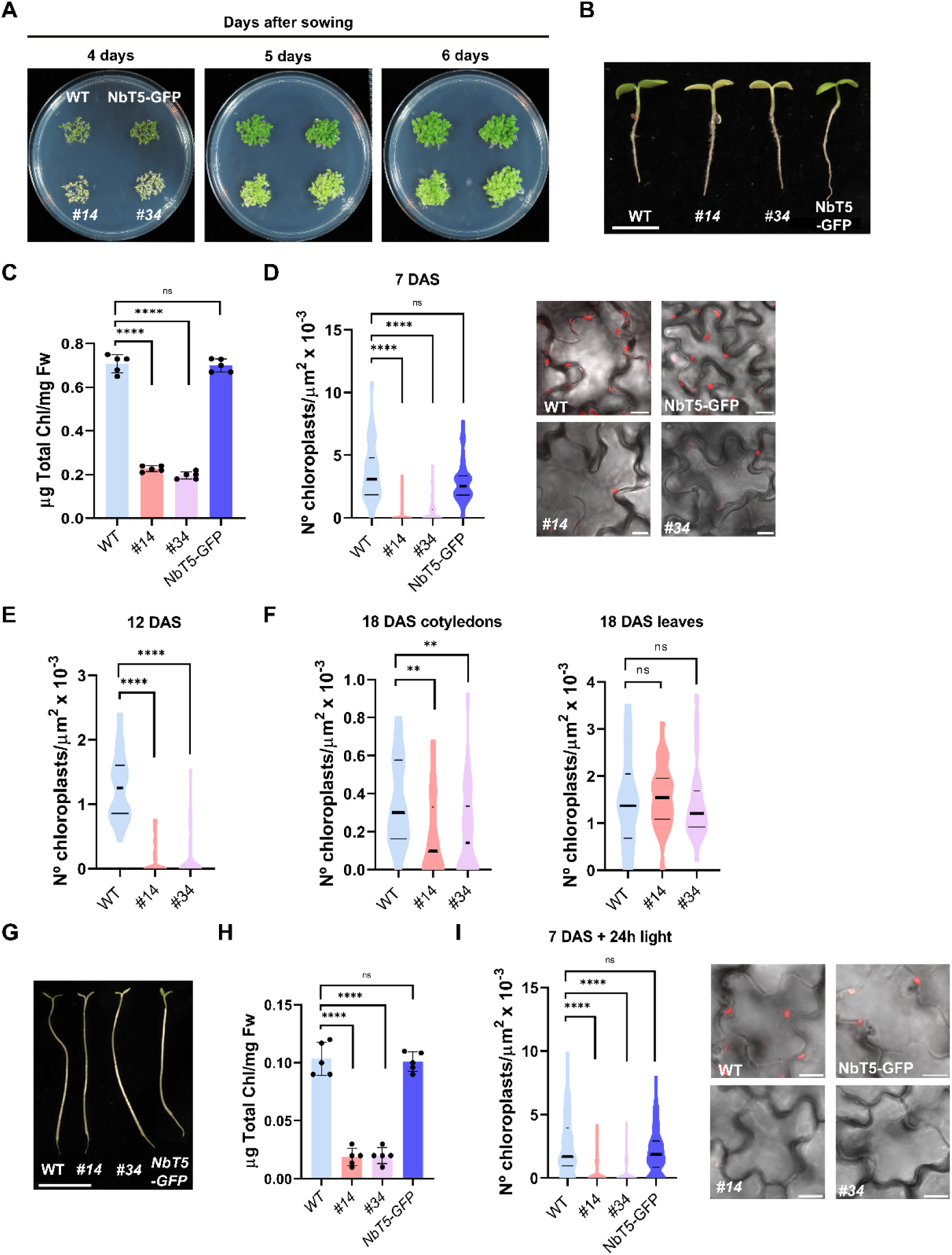
Loss of *NTMC2T5* impairs cotyledon greening and chloroplast development during early seedling development and de-etiolation. (A-F) Seedlings were grown on sucrose-free MS medium under long-day conditions. **(A)** Top-view photographs of WT, *nbt5* mutants (lines *#14* and *#34*), and complemented NbNTMC2T5-GFP (NbT5-GFP) lines at different days after sowing, highlighting the cotyledon color differences among genotypes. **(B)** Side-view photographs of WT, *nbt5* mutants (lines *#14* and *#34*), and complemented NbT5-GFP lines at 5 days after sowing. Scale bar: 0.5 cm. **(C)** Total chlorophyll content at 7 days after sowing. Bars represent the mean ± SD. Data were analyzed by one-way ANOVA followed by Dunnett’s multiple comparison test. **(D)** Chloroplast density in cotyledon epidermal cells of 7-day-old seedlings (left panel). Chloroplast density was calculated for individual cells by dividing the total number of chloroplasts within each cell by its measured cell area (chloroplasts µm⁻²). Representative single-plane confocal images showing chloroplasts (red) are shown in the right panel. Cell outlines were visualized by bright-field microscopy and chlorophyll autofluorescence was detected using 633 nm excitation. Scale bar: 10 µm. **(E)** Chloroplast density in cotyledon epidermal cells of 12-day-old seedlings, calculated as in (D). **(F)** Chloroplast density in cotyledon and leaf epidermal cells of 18-day-old seedlings, calculated as in (D). **(G–I)** Seedlings were germinated and grown for 7 days in darkness on sucrose-free MS medium, then transferred to long-day conditions for 24 h before analysis. **(G)** Photographs of de-etiolated WT, *nbt5* mutants (lines *#14* and *#34*), and complemented NbT5-GFP line. Scale bar: 1 cm. **(H)** Total chlorophyll content after de-etiolation. Bars represent the mean ± SD. Data were analyzed by one-way ANOVA followed by Dunnett’s multiple comparison test. **(I)** Chloroplast density in cotyledon epidermal cells after de-etiolation (left panel), as calculated in (D). Representative single-plane confocal images showing chloroplasts (red) in each genotype are shown in the right panel. Cell outlines were visualized by bright-field microscopy and chlorophyll autofluorescence was detected using 633 nm excitation. Scale bar: 10 µm. **(D–F, I)** Data are shown as violin plots with the median and quartiles indicated. Data were analyzed using the Kruskal-Wallis test followed by Dunn’s multiple comparison test. **(C–F, H, I)** Asterisks indicate statistically significant differences (ns, not significant; **, *P* ≤ 0.01; ****, *P* ≤ 0.0001). *Abbreviations:* DAS, days after sowing; NbT5, NbNTMC2T5.

To determine whether the requirement for *NbT5* extends to other developmental transitions involving *de novo* chloroplast formation, we examined seedling de-etiolation. During skotomorphogenic growth, proplastids differentiate into etioplasts, which rapidly convert into chloroplasts upon illumination through extensive membrane biogenesis and thylakoid assembly (Pipitone *et al*., 2021). Seedlings were germinated in darkness in the absence of exogenous sucrose and exposed to light after 6–7 days. Following illumination, WT and complemented seedlings underwent normal photomorphogenic development, characterized by cotyledon greening and apical hook opening (Fig. 3G) (Wang *et al*., 2020). In contrast, *nbt5* mutants failed to green and displayed a pronounced albino phenotype. Consistent with this phenotype, total chlorophyll content, as well as chlorophyll *a* and chlorophyll *b* levels, were dramatically reduced relative to WT and complemented seedlings (Fig. 3H and Fig. S4G). Additionally, confocal microscopy revealed a more striking defect. Chloroplast autofluorescence was nearly undetectable in *nbt5* cotyledons with or without exogenous source of sucrose (Fig. 3I and S4H). Quantification confirmed an almost complete absence of chlorophyll autofluorescence-positive structures in mutant cotyledon cells. Together, these findings indicate that NTMC2T5 is required for efficient chloroplast development during early seedling establishment and de-etiolation, but becomes largely dispensable for chloroplast maintenance in mature leaves.

### NTMC2T5 is required for plastid differentiation and internal membrane organization

The severe reduction in chlorophyll autofluorescence-positive structures observed in *nbt5* cotyledons under both light-grown and de-etiolation conditions prompted us to examine how loss of NTMC2T5 affects plastid differentiation. We therefore analyzed plastid ultrastructure in cotyledon epidermal cells by transmission electron microscopy (TEM) following de-etiolation. In WT seedlings, plastids representing different developmental states were readily identified (Fig. 4A,C,E). Etioplasts containing well-defined prolamellar bodies were frequently observed (Fig. 4C, empty white arrowheads) together with chloroplasts exhibiting organized internal membrane systems (Fig. 4E). Electron-dense plastoglobule-like structures were also observed (Fig. 4C, white filled arrow). In contrast, plastids in *nbt5 #14* seedlings displayed markedly altered ultrastructure (Fig. 4B,D,F). These plastids lacked recognizable prolamellar bodies, thylakoid membranes, and organized thylakoid membrane systems and instead contained disorganized vesicular structures and circular membrane profiles resembling aberrant prothylakoid-like membranes (Fig. S5A,B).

**Figure 4.**
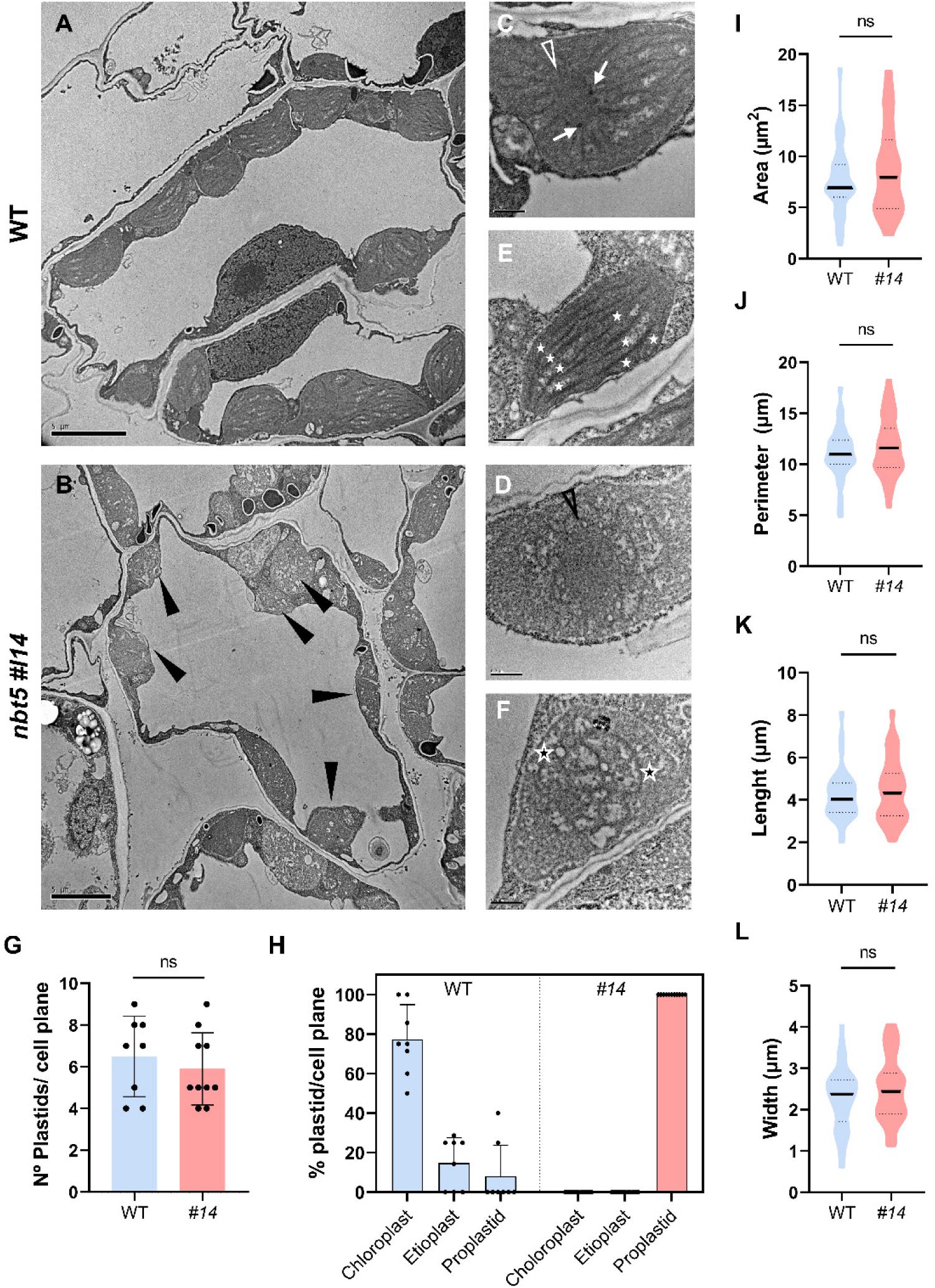
Loss of *NTMC2T5* disrupts the internal organization of plastids. Transmission electron micrographs of *Nicotiana benthamiana* cotyledons from WT **(A, C and E)** and *nbt5 #14* **(B, D and F)** plants grown for 7 days in darkness followed by 1 day in light. **(A, B)** Overview of cotyledon cells in WT (A) and *Nbt5* (B). **(C, E)** WT plastids predominantly included etioplasts (C) and young chloroplasts (E). **(D, F)** In *nbt5* cotyledons, no young chloroplasts were observed. Instead, abnormal etioplast-like structures **(D)** and proplastids-like structures **(F)** were present. **(A–F)** Scale bars: 5 µm (A, B) and 500 nm (C–F). **(G)** Total number of plastids per cell plane. Bars represent the mean ± SD, with each dot representing one cell section. **(H)** Distribution of plastid types, calculated as the percentage of each plastid type per cell plane. Bars represent the mean ± SD, with each dot representing one cell section. **(I–L)** Plastid size parameters measured from the micrographs in WT and *nbt5* plants: area **(I)**, perimeter **(J)**, length **(K)** and width **(L)**. Data are shown as violin plots with the median and quartiles indicated. **(G, I–L)** Data were analyzed by unpaired two-tailed Student’s *t*-test (ns, not significant, *P* > 0.05). *Symbols in TEM micrographs*: developing grana (white stars), lamellar prothylakoids within proplastid-like structures (black stars), plastoglobules (white filled arrows), putative remnants of the prolamellar body (empty black arrowheads), remnants of the prolamellar body (empty white arrowheads). *Abbreviations: nbt5, nbntmc2t5*.

We next classified plastids according to their ultrastructural characteristics. Plastid number per cell plane did not differ significantly between WT and *nbt5 #14* cotyledons (Fig. 4G), whereas the distribution of plastid developmental classes was markedly different (Fig. 4H). In WT seedlings, approximately 77% of plastids were classified as chloroplasts, 15% as etioplasts, and 8% as proplastids. By contrast, plastids in *nbt5 #14* seedlings lacked the defining ultrastructural features of etioplasts and chloroplasts and were classified as undifferentiated proplastid-like plastids. We then asked whether the altered plastid differentiation was accompanied by changes in plastid size. In higher plants, proplastids are typically smaller than mature chloroplasts, which undergo substantial expansion during differentiation (Pyke and Leech, 1992; Robertson *et al*., 1995; Yadav *et al*., 2019). Measurements of plastid area, perimeter, length and width revealed no significant differences between WT and *nbt5 #14* plastids (Fig. 4I–L). Thus, loss of NTMC2T5 was not associated with a detectable reduction in plastid size but was accompanied by a pronounced alteration in internal membrane organization.

Collectively, these data indicate that NTMC2T5 is required for the acquisition of the ultrastructural features characteristic of differentiated plastids. In particular, the absence of recognizable prolamellar bodies and organized internal membrane systems in *nbt5 #14* plastids suggests that NTMC2T5 is important for the membrane remodeling associated with early plastid differentiation. These findings are consistent with a role for NTMC2T5 early in plastid differentiation, before the establishment of mature chloroplast architecture.

### NTMC2T5 proteins are enriched at ER–chloroplast interfaces

Members of the SMP-domain protein family are typically associated with membrane contact sites, where they can function as membrane tethers and/or lipid transfer proteins (Toulmay and Prinz, 2012; AhYoung *et al*., 2015; Reinisch and De Camilli, 2016; Jeyasimman and Saheki, 2020). To investigate whether NTMC2T5 proteins are associated with ER–chloroplast interfaces, we first examined the subcellular localization of NbT5-GFP in the complemented *N. benthamiana* line. Confocal imaging revealed a punctate distribution of NbT5-GFP, with fluorescence enriched at discrete regions adjacent to chloroplasts (Fig. 5A). This pattern suggested that NbT5 is enriched at discrete membrane-associated regions around chloroplasts.

**Figure 5.**
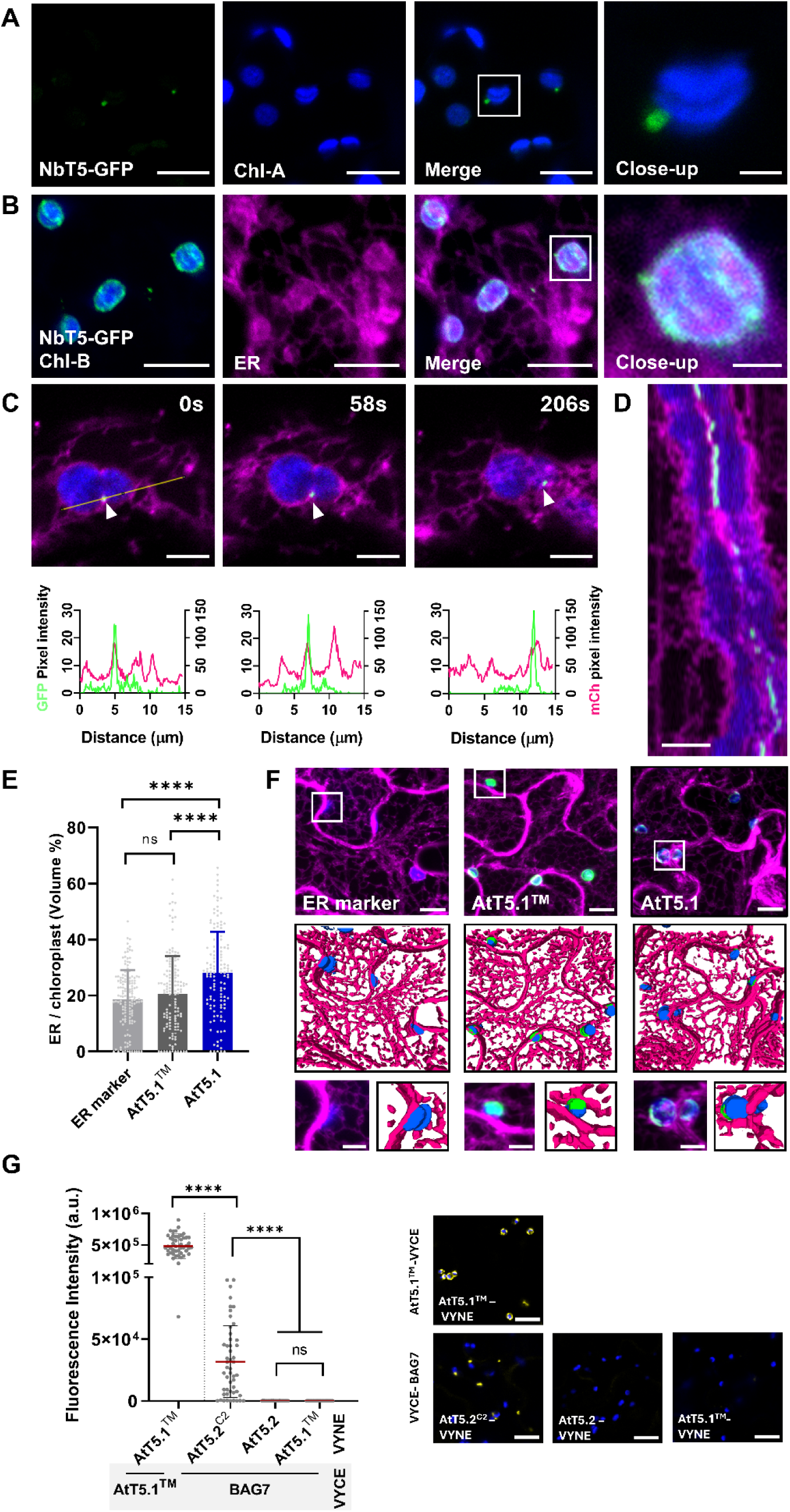
NTMC2T5 proteins are located at ER–chloroplast interfaces. **(A)** Confocal images (single plane) of a stable *N. benthamiana* line expressing NbT5-GFP. Images of the individual channels as well as the merged images are shown. Boxed region is magnified in the insets. Scale bar: 10 µm; inset scale bar: 2 µm. **(B)** Maximum-intensity Z-projections of confocal images of *N. benthamiana* leaves transiently co-expressing NbT5-GFP and ER marker fused to mCh (ER-mCh). NbT5-GFP partially co-localized with the ER signal. The boxed region is magnified in the inset. Individual channels and the merged image are shown. Scale bar: 10 µm; inset scale bar: 5 µm. **(C)** Time-lapse confocal images of epidermal cells of the stable NbT5-GFP line transiently expressing ER-mCh. Representative frames of the time series are shown. Fluorescence intensity profiles measured along the indicated line are displayed below the corresponding images; the same line position (shown in the first frame) was used for all time points. Scale bar: 5 µm. **(D)** Kymograph generated from the time-lapse series shown in (C), corresponding to the line indicated in the first frame, illustrating the temporal dynamics of the NbT5-GFP and ER-mCh signals along the selected region. Scale bar: 5 µm. **(E)** Quantification of the ER volume surrounding chloroplasts in *N. benthamiana* leaves expressing ER-mCh alone or co-expressing AtT5.1^TM^-GFP or AtT5.1-GFP. ER volume was calculated as described in Methods. Bars represent the mean ± SD. Data were analyzed using the Kruskal-Wallis test followed by Dunn’s multiple comparison test (*P* < 0.05). **(F)** Representative confocal Z-stacks and 3D reconstructions of the cells quantified in (E). Boxed regions are magnified in the insets. Corresponding magnified 3D reconstructions are also shown. Scale bar: 10 µm; inset scale bar: 5 µm. **(G)** BiFC assay in *N. benthamiana* epidermal cells testing the interaction between AtT5.2 variants and the ER-localized protein BAG7. Quantification of the Venus signal is shown in the left panel and representative single-plane confocal images in the right panel. Leaves were transiently co-transformed with VYCE-BAG7 and a truncated version of AtT5.2 ending at the C2 domain (AtT5.2^C2^, M1–D549), full-length AtT5.2 or AtT5.1^TM^ (M1–G160; negative control), each fused to VYNE. AtT5.1^TM^-VYNE co-expressed with AtT5.1^TM^-VYCE served as a positive control. Reconstituted Venus fluorescence was detected at discrete sites on the chloroplast surface only with AtT5.2^C2^-VYNE. Fluorescence intensities (a.u.) were measured in ImageJ from randomly selected regions of interest in non-saturated images. Data were analyzed using the Kruskal-Wallis test followed by Dunn’s multiple comparison test. Note that the images shown are saturated and are intended for qualitative comparison only. Scale bar: 20 µm. **(A-D, F, G)** Chloroplasts were visualized by chlorophyll autofluorescence (blue). GFP fluorescence (A–D, F) is shown in green and reconstituted YFP fluorescence (G) in yellow. **(B–D, F)** ER-mCh is shown in magenta. *Abbreviations:* AtT5.1, AtNTMC2T5.1; AtT5.2, AtNTMC2T5.2; NbT5, NbNTMC2T5; TM, transmembrane region.

We next examined the spatial relationship between NbT5 and the ER. Co-expression of NbT5-GFP with the ER marker CD3-959 (Nelson et al., 2007) revealed extensive spatial overlap of both signals in regions surrounding chloroplasts (Fig. 5B). High-magnification views further showed coincident enrichment of NbT5 and ER signals at discrete regions adjacent to the chloroplast perimeter. These observations are consistent with an association of NbT5 with ER membranes at chloroplast-associated regions. Similarly, co-expression of the ER marker with its *Arabidopsis* homolog AtT5.1 resulted in extensive overlap of the two fluorescent signals around chloroplasts (Fig. S6), further suggesting that NTMC2T5 proteins may associate with ER membranes.

To determine whether the association of NbT5 with the ER was maintained over time, we performed live-cell imaging in complemented *N. benthamiana* plants expressing NbT5 together with the ER marker. Time-lapse imaging revealed persistent enrichment of NbT5 at ER regions surrounding chloroplasts over the recording period. This spatial association was maintained despite substantial ER remodeling and chloroplast movement (Fig. 5C), indicating that the enrichment of NbT5 at chloroplast-associated ER regions persists during dynamic changes in the cortical membrane system. We further generated a kymograph along a representative line scan spanning an ER–chloroplast interface (Fig. 5D). The resulting kymograph showed sustained co-migration of NbT5 and the ER marker along the same spatial coordinates over time. Together, these observations support a persistent association of NbT5 with ER membranes at chloroplast-associated regions.

We next asked whether NTMC2T5 proteins could influence the extent of ER– chloroplast membrane apposition. If AtT5.1 contributes to the association between the ER and chloroplasts, its expression would be expected to increase the extent of ER signal overlapping with chloroplasts. To test this possibility, we transiently expressed an ER marker alone or together with AtT5.1-GFP or the chloroplast-targeted transmembrane segment AtT5.1^TM^-GFP in *N. benthamiana* leaves. Three-dimensional reconstructions obtained from multiple focal planes were used to quantify ER– chloroplast fluorescence overlap around individual chloroplasts. Expression of full-length AtT5.1 significantly increased ER–chloroplast overlap compared with expression of the ER marker alone or AtT5.1^TM^ (Fig. 5E). Representative three-dimensional reconstructions further illustrated the increased extent of ER signal surrounding chloroplasts in cells expressing full-length AtT5.1 (Fig. 5F). These results indicate that AtT5.1 can increase ER–chloroplast membrane apposition and suggest that sequences outside the chloroplast-targeting transmembrane region contribute to this activity.

To further assess the proximity of NTMC2T5 proteins to the ER, we performed bimolecular fluorescence complementation (BiFC) using the ER-resident protein BAG7. Whereas AtT5.1^TM^-VYNE did not reconstitute Venus fluorescence when co-expressed with VYCE-BAG7, the AtT5.2^C2-^VYNE construct (M1–D549) produced a clear BiFC signal at discrete regions surrounding chloroplasts (Fig. 5G), consistent with close molecular proximity between AtT5.2^C2^ and an ER-resident protein. Importantly, the absence of signal with AtT5.1^TM^-VYNE was not due to loss of VYNE functionality, as strong fluorescence was observed when AtT5.1^TM^-VYNE was co-expressed with AtT5.1^TM^-VYCE. Unexpectedly, full-length AtT5.2-VYNE did not produce detectable BiFC with VYCE-BAG7. Because NTMC2T5 proteins contain a predicted disordered C-terminal region followed by a short hydrophobic segment, the C-terminal VYNE fusion may affect the accessibility or orientation of the split-Venus fragment. Together, these results support a close association of NTMC2T5 with the ER and are consistent with its enrichment at ER–chloroplast interfaces.

### NTMC2T5 is required for proper lipid remodeling during plastid differentiation

The severe ultrastructural defects observed in *nbt5* plastids, which fail to establish normal prolamellar bodies and thylakoid membranes despite attaining near-normal organelle size, suggested that NTMC2T5 is required for the establishment of normal plastid internal membrane systems. Because SMP-domain proteins typically mediate lipid exchange at membrane contact sites, we investigated whether the loss of NTMC2T5 affects lipid homeostasis during plastid development. As an initial approach, we examined seed germination and lipid composition in dry seeds. Germination was unaffected by the mutation, reaching close to 100% in both WT and *nbt5* seeds three days after sowing (Fig. S7A). Consistent with the absence of developmental defects at this stage, lipidomic analyses revealed no significant differences in total lipid content, fatty acid composition, or the abundance of major polar and neutral lipid classes between WT and mutant seeds (Fig. S7B–E). These results suggest that NTMC2T5 is dispensable for lipid accumulation and storage during seed maturation.

We next asked whether NTMC2T5 becomes important during the transition from proplastids into photosynthetically competent plastids. During skotomorphogenesis and subsequent de-etiolation, plastids undergo extensive membrane biogenesis, requiring the coordinated synthesis and exchange of lipids between the ER and plastid envelopes (Pipitone *et al*., 2021). Given the association of NTMC2T5 with ER–chloroplast interfaces, we hypothesized that its function may become particularly important during this developmental transition. Lipidomic profiling of de-etiolated seedlings revealed no significant differences in total lipid content between WT and *nbt5* lines (Fig. 6A), indicating that overall lipid accumulation was not impaired. In contrast, pronounced changes were detected in fatty acid composition (Fig. 6B). Both independent mutant lines accumulated higher proportions of C18:1 and C18:2 fatty acids, whereas C18:3 was strongly reduced relative to WT seedlings. Together, these changes are indicative of reduced enrichment in polyunsaturated fatty acids, which are characteristic of photosynthetic membranes (McConn and Browse, 1998).

**Figure 6.**
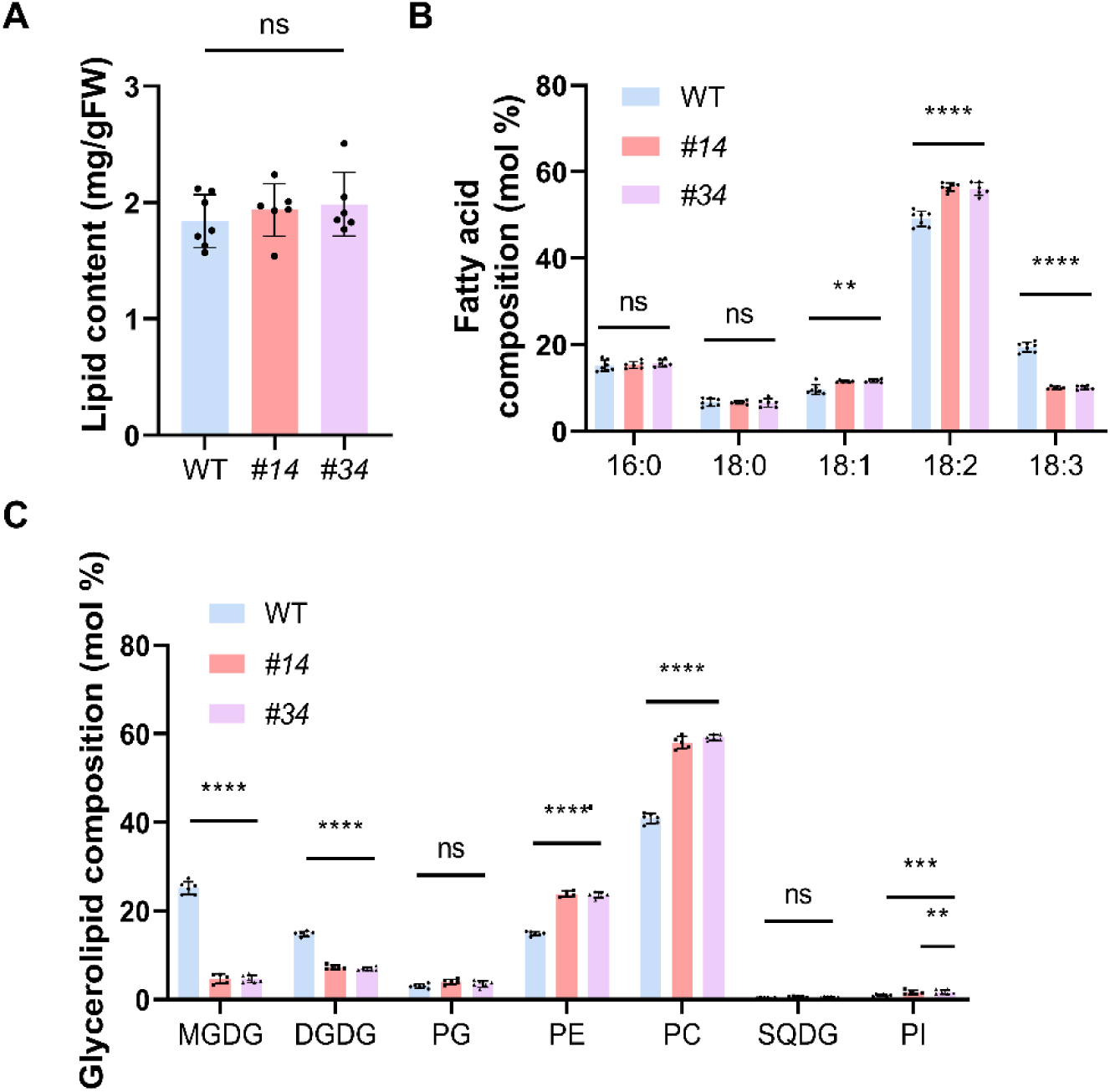
*ntmc2t5* mutants show altered lipid composition during de-etiolation. Seedlings of WT and *nbt5* lines *#14* and *#34* were germinated and grown in darkness for 6 days, followed by 24 h under long-day conditions. Total lipids were extracted and analyzed by mass spectrometry. **(A)** Total lipid content of seedlings (mg g⁻¹ fresh weight). (**B)** Fatty acid distribution (mol %). **(C)** Glycerolipid composition (mol %). **(A–C)** Bars represent the mean ± SD. Data were analyzed by one-way ANOVA (A) or two-way ANOVA (B, C) followed by Dunnett’s multiple comparison test relative to WT. Asterisks indicate significant differences (ns, not significant; **, *P* ≤ 0.01; ****, *P* ≤ 0.0001). *Abbreviations:* DGDG, digalactosyldiacylglycerol; FW, fresh weight; MGDG, monogalactosyldiacylglycerol; NbT5, NbNTMC2T5; PC, phosphatidylcholine; PE, phosphatidylethanolamine; PG, phosphatidylglycerol; PI, phosphatidylinositol; SQDG, sulfoquinovosyldiacylglycerol.

Analysis of glycerolipid classes revealed more pronounced changes in lipid partitioning (Fig. 6C). The major plastid galactolipids, monogalactosyldiacylglycerol (MGDG) and digalactosyldiacylglycerol (DGDG), were strongly reduced in *nbt5* seedlings with MGDG decreasing by approximately 80%, and DGDG by nearly 50% relative to WT. In contrast, phospholipids typically associated with extraplastidial membranes accumulated to significantly higher levels in the mutant. Phosphatidylethanolamine (PE), phosphatidylcholine (PC), and phosphatidylinositol (PI) all increased relative to WT seedlings, while phosphatidylglycerol (PG) remained largely unchanged (Fig. 6C). Consequently, loss of NTMC2T5 results in a pronounced shift in glycerolipid composition, characterized by depletion of plastid galactolipids and enrichment of ER-derived phospholipids. Importantly, these changes coincided with the developmental stage at which *nbt5* plastids fail to establish normal internal membrane systems. Given that galactolipids constitute the major structural lipids of prolamellar bodies and thylakoid membranes, the strong reduction in MGDG and DGDG is consistent with impaired plastid membrane biogenesis in the mutant. Together, these results demonstrate that NTMC2T5 is not required for lipid storage during seed development but becomes essential during plastid differentiation. The loss of NTMC2T5 causes a profound imbalance between plastid galactolipids and extraplastidial phospholipids during de-etiolation, providing a mechanistic explanation for the failure of *nbt5* plastids to assemble normal internal membrane structures.

## DISCUSSION

Membrane contact sites are increasingly recognized as important platforms for coordinating lipid exchange and organelle function (Andersson et al., 2007; Yao et al., 2023; Renna et al., 2024; Huercano et al., 2025, 2026). Plastid membrane biogenesis depends on coordinated lipid metabolism within plastids and on the import of ER-derived lipid precursors (Hölzl and Dörmann, 2019; Pipitone et al., 2021). Several systems have been implicated in lipid trafficking at ER–chloroplast contact sites, including the TGD complex, SFH5 and SFH7, and the VAP27–ORP2A complex, (Wang *et al*., 2012a; Fan *et al*., 2015; Yao *et al*., 2023; Renna *et al*., 2024). Nevertheless, the molecular components and developmental roles of these interfaces remain incompletely understood. Here, we identify NTMC2T5 as a previously uncharacterized SMP-domain protein family associated with the chloroplast envelope and provide genetic, cell biological, ultrastructural, and lipidomic evidence linking NTMC2T5 to lipid homeostasis and plastid differentiation.

Our sequence similarity analysis places NTMC2T5 in a distinct SMP-domain clade, separate from the canonical plant SYT1–5 proteins, which cluster with their metazoan homologs (Fig. 1B). NTMC2T5 homologs were detected across all major Archaeplastida lineages examined, from red algae and glaucophytes to green algae and land plants (Fig. 1D). This distribution is most consistent with an ancient origin followed by lineage-specific losses, suggesting that NTMC2T5 was already present in the last common ancestor of Archaeplastida. Notably, the complete TM–SMP–C2–HR architecture was recovered in every lineage in which homologs were detected, indicating that this domain organization has been maintained since the origin of Archaeplastida. This conservation prompted us to ask how each region contributes to the targeting and activity of these proteins. NTMC2T5 proteins contains an N-terminal membrane-targeting region, an SMP domain, a C2 domain and a short C-terminal hydrophobic region (Fig. 2A). The N-terminal region was sufficient to target GFP to the chloroplast periphery (Fig. 2D and S2A), whereas its removal redirected the protein to an ER-like reticulate pattern (Fig. S2B,C), identifying this region as a major determinant of chloroplast-envelope localization. However, full-length AtT5.1 protein had functions beyond targeting: it promoted perinuclear chloroplast accumulation (Fig. 2E), and increased ER–chloroplast fluorescence overlap, effects that were not reproduced by the isolated targeting region (Fig. S6). Thus, domains outside the membrane anchor are likely to contribute to the organization or activity of NTMC2T5 at the chloroplast surface. Consistent with this, the full-length protein displayed a punctate distribution at the chloroplast surface, whereas the isolated targeting region distributed uniformly around chloroplasts (Fig. 2C,D), suggesting that NTMC2T5 concentrates at specific regions of the envelope.

Several observations indicate that these regions correspond to specialized chloroplast membrane domains associated with the ER. Full-length proteins displayed a punctate distribution at the chloroplast surface (Fig. 2C, 5A), with enrichment at regions adjacent to the ER (Fig. 5B,C) and this distribution persisted during ER remodeling and chloroplast movement (Fig. 5C). Recent *in situ* imaging has highlighted the dynamic nature of chloroplast positioning in intact leaves (Geng *et al*., 2025), emphasizing that these contacts occur within a highly dynamic cellular environment. Independent proteomic and imaging evidence also points to functionally distinct membrane domains within the chloroplast. LaBrant *et al*. (2026) identified a chloroplast membrane fraction intermediate between the envelope and thylakoid membranes that was enriched in lipid-metabolic proteins and membrane-organization factors, including NTMC2T5.1, which exhibited discrete punctate localization (LaBrant *et al*., 2026). BiFC between the C2-containing region of AtT5.2 and the ER-resident protein BAG7 further supported close molecular proximity between NTMC2T5 and the ER (Fig. 5G). Moreover, full-length AtT5.1 increased ER–chloroplast fluorescence overlap, whereas its isolated targeting region did not (Fig. 5E,F), indicating that regions outside the membrane anchor contribute to the spatial organization of the two membranes.

The developmental phenotype of *nbt5* knock-out mutant lines (Fig.3 and Fig. 4) supports that NTMC2T5 activity is particularly important during early plastid differentiation, when the lipid requirements of the nascent plastid are high and its capacity for autonomous membrane-lipid production is not yet fully established. Loss of *NbT5* caused pale or variegated cotyledons (Fig. 3A,B,G), reduced chlorophyll accumulation (Fig.3C,H) and a strong reduction in chlorophyll autofluorescence-positive structures (Fig. 3D,E,I) during seedling development, whereas complementation restored these defects (Fig. 3). The requirement for NTMC2T5 was markedly reduced in mature leaves, (Fig. S4D-F). This developmental specificity may also reflect differences in plastid ontogeny between organs. In cotyledons, etioplasts derive from embryo mesophyll proplastids and chloroplast development depends strongly on *de novo* membrane assembly, whereas leaf chloroplasts arise from meristematic proplastids and subsequently proliferate mainly through division (Waters and Langdale, 2009; Pogson and Albrecht, 2011). This temporal phenotype is consistent with a role in *de novo* plastid membrane biogenesis rather than in the maintenance of established chloroplasts.

At the ultrastructural level, the defect was more precisely localized to the internal membranes of the plastid. Plastids in *nbt5* lines retained near-normal number and size but lacked recognizable prolamellar bodies and organized thylakoid membranes and instead contained disorganized vesicular and circular membrane structures (Fig. 4 and S5). Thus, loss of NTMC2T5 does not appear to primarily impair plastid formation or expansion but rather the establishment of the internal membrane architecture characteristic of differentiated etioplasts and chloroplasts. Because prolamellar-body organization depends strongly on galactolipid composition, particularly the relative abundance of MGDG and DGDG (Jarvis *et al*., 2000; Fujii *et al*., 2017, 2018, 2019a,*b*) the ultrastructural phenotype provides a direct cellular context for the lipidomic changes observed in the mutant.

Lipidomic analysis revealed that NTMC2T5 is dispensable for bulk lipid accumulation in dry seeds (Fig. S7) but becomes important during de-etiolation (Fig. 6). Total lipid abundance was not significantly altered in mutant seedlings (Fig. 6A), whereas lipid composition changed markedly: MGDG and DGDG decreased, whereas PC, PE and PI increased (Fig. 6C). Fatty-acid composition was also shifted, with increased C18:1 and C18:2 and a strong reduction in C18:3 (Fig 6B). The coordinated depletion of plastid galactolipids and accumulation of phospholipids is therefore more consistent with altered lipid partitioning, delivery, or remodeling than with a general reduction in lipid synthesis. Importantly, these changes occurred at the developmental stage when mutant plastids failed to establish normal internal membrane systems, providing a biochemical correlate of the ultrastructural phenotype.

Because plastids are not part of the vesicular trafficking system, lipid exchange between the ER and plastids is thought to depend predominantly on non-vesicular transport at ER–plastid contact sites (Wang and Benning, 2012; Hurlock *et al*., 2014; Block and Jouhet, 2015; LaBrant *et al*., 2018; Michaud and Jouhet, 2019; Huercano *et al*., 2025, 2026). The presence of an SMP domain provides a plausible molecular basis for a role in lipid handling at ER-plastid interfaces. SMP domains are established lipid-transfer modules (Schauder *et al*., 2014; AhYoung *et al*., 2015; Reinisch and De Camilli, 2016; Jeyasimman and Saheki, 2020; Ruiz-Lopez *et al*., 2021; Wang *et al*., 2023). Our data therefore support a model in which NTMC2T5 contributes to ER–plastid lipid homeostasis during early plastid differentiation, but they do not distinguish between direct role in lipid transfer and an indirect role in organizing membrane interfaces or recruiting other lipid-trafficking components.

We therefore propose that NTMC2T5 promotes efficient lipid homeostasis at the ER–plastid interface during the early stages of plastid differentiation. Reduced NTMC2T5 activity would impair the delivery or partitioning of lipid precursors required for galactolipid-rich internal membrane formation, resulting in reduced MGDG and DGDG and failure to establish normal prolamellar bodies and thylakoid membranes. Once chloroplasts have matured, endogenous plastid lipid synthesis and/or alternative lipid-exchange pathways may partially compensate for the loss of NTMC2T5, accounting for the strong developmental specificity of the phenotype. Direct lipid-flux measurements, biochemical assays of SMP-dependent lipid transfer, and domain-specific mutant analyses will be required to determine whether NTMC2T5 directly transfers particular lipid species or primarily organizes the membrane environment in which lipid exchange occurs.

Together, our findings identify NTMC2T5 as an Archaeplastida-conserved SMP-domain protein family required for efficient plastid differentiation and establish a connection between NTMC2T5, ER-associated chloroplast membrane regions, and lipid remodeling during chloroplast biogenesis. More broadly, these results suggest that specialized membrane interfaces may contribute not only to lipid exchange between organelles but also to the developmental establishment of chloroplast membrane identity.

## MATERIALS AND METHODS

### Plant Material and growth conditions

*N. benthamiana* seeds were surface-sterilized and sown on half-strength Murashige and Skoog (MS) agar medium (0.8% agar). The plates were placed either horizontally or vertically in a growth chamber under cool-white light (120 μmol photons m^−2^ s^−1^) with controlled environmental conditions (25 °C, 16-h light/8-h dark cycle), unless specified otherwise. When necessary, seedlings were transferred to soil composed of an organic substrate and vermiculite in a 4:1 (v/v) ratio and grown under the same temperature and photoperiod conditions. Freshly harvested seeds were used for all experiments.

### Identification and CLANS-based clustering of SMP-domain proteins in *Arabidopsis thaliana* and *Nicotiana benthamiana*

To identify SMP domain-containing proteins in plants, iterative homology searches were initiated using the SMP domain of human extended Synaptotagmin 1 (HsE-Syt1; residues D135–V313) as the initial query. Profile-based searches were performed with HHpred (Zimmermann *et al*., 2018) against the *A. thaliana* proteome (TAIR10; accessed 20 June 2017; database: PDB_mmCIF70), and hits with probability scores ≥50% were retained. Newly identified SMP-containing proteins were subsequently used as queries in additional rounds of searching until no further candidates were recovered. To identify SMP-domain proteins in *N. benthamiana*, PSI-BLAST searches were performed against the predicted proteome available at the Sol Genomics Network (*N. benthamiana* v1.0.1). Candidate proteins were analyzed with InterProScan (Blum *et al*., 2025) to determine their domain architecture, and only proteins containing a predicted SMP domain were retained for subsequent analyses.

The SMP proteins identified in this study were analyzed together with previously reported SMP-containing proteins from *H. sapiens*, *S. cerevisiae* and several plant species described by Craxton (Craxton, 2007). Sequence similarity relationships were analyzed using CLANS (Frickey and Lupas, 2004), which clusters sequences based on pairwise similarity scores. Pairwise similarities were calculated using the BLOSUM62 scoring matrix and an E-value cutoff of 1 × 10⁻⁹. Clustering was visualized using Java implementation of CLANS with default settings. Only connections with *P*-values below 0.1 were considered for downstream interpretation. All sequences used for CLANS analysis are provided in Supplementary File 1.

### Identification and phylogenetic analysis of *N. benthamiana* NTMC2T5 homologs across Archaeplastida

The NbT5 protein sequence was used to identify putative homologs across representative species spanning the major Archaeplastida lineages. PSI-BLAST searches were carried out against the NCBI non-redundant (nr), RefSeq Protein, clustered nr, and TSA_nr (Transcriptome Shotgun Assembly) databases using default parameters to maximize sequence recovery from both annotated genomes and transcriptome-derived datasets. Representative species were selected to maximize taxonomic coverage. When no significant hits were recovered from a given species, additional PSI-BLAST searches restricted by the corresponding taxonomic identifier (TaxID) were performed to interrogate all available species within that clade. Candidate proteins were analyzed with InterProScan (Blum *et al*., 2025) and TOPCONS (Tsirigos *et al*., 2015) to determine their domain architecture. Only proteins containing the combination of TM, SMP, C2, and hydrophobic domains identified in *Arabidopsis* and *Nicotiana* NTMC2T5 proteins were retained for further analysis.

Then, protein sequences were aligned using MAFFT v7 (Katoh *et al*., 2019) with default settings. Phylogenetic relationships were inferred by the Maximum Likelihood method implemented in MEGA12 (Kumar *et al*., 2024). The best-fitting amino acid substitution model was selected by model testing, and the final tree was reconstructed using the LG+G+I+F model. Rate heterogeneity among sites was modeled using a discrete Gamma distribution with five categories (α = 1.9437), and 0.84% of sites were treated as evolutionarily invariant. Node support was estimated from 500 bootstrap replicates, and branches with bootstrap values <50% were collapsed. All other parameters were left at their default settings.

### Proteins localization and domain prediction

Subcellular localization of SMP proteins was predicted using DeepLoc 2.1 (Ødum *et al*., 2024) predictor server. Protein domains were predicted using InterPro (Blum *et al*., 2025), except for transmembrane domains which were predicted using TOPCONS (Tsirigos *et al*., 2015). Three-dimensional structures were predicted using AlphaFold3 (Abramson *et al*., 2024).

### Single guide RNA design

The online tools CRISPOR (http://crispor.tefor.net) (Concordet and Haeussler, 2018) and CCTop (https://cctop.cos.uni-heidelberg.de) (Stemmer *et al*., 2015) were used to select CRISPR targets. Results from both tools were compared to select guides with high efficacy (CCTop score >0.74; CRISPOR Mor-Mateos score >60). Targets with no predicted off-targets or a low number of off-targets were prioritized. When off-targets were predicted, guides with a higher number of mismatches in the seed sequence were selected. Predicted off-target sites are listed in Supplemental Table S3.

### Plasmid constructs/Cloning

#### Gateway-based cloning

The coding DNA sequences (CDS) without stop codons of *AtT5.1* (*AT1G50260*)*, AtT5.2 (AT3G19830)* and *AtBAG7* (AT5G62390) were PCR-amplified from Arabidopsis Col-0 cDNA, using the primers listed in Table S2 and cloned into pENTR/Zeo vector by BP cloning kit (Invitrogen). CDS of *NbT5* (*NIBEN101SCF08519G00002*) was PCR-amplified from *N. benthamiana* cDNA and cloned as described above. The truncated versions AtT5.1^TM^ (M1 to G160), AtT5.1^ΔTM^ (E183-P675), AtT5.2^TM^ (M1-G179), AtT5.2^C2^ (M1-D549) and NbT5^TM^ (M1-L185) were PCR amplified using the primers detailed in Supplemental Table S2 and built into pENTR/Zeo vectors. All pENTR/Zeo vectors were verified by sequencing. The final Gateway-expression vector was constructed by LR-reaction (Invitrogen). The pENTR vectors, pEN-L4-pUBQ10-R1 and pENT L2-GFP-R3 were used with the pDEST vector pH7m34GW. The ER marker (CD3-959, ER-mCh) was retrieved from (Nelson *et al*., 2007). For BiFC experiments, the pENTR/Zeo vectors which included the coding regions of AtT5.2, AtT5.2^C2^, AtT5.1^TM^, or BAG7 were transferred into Gateway-compatible binary vectors: pDEST-^GW^VYNE, pDEST-^GW^VYCE or pDEST-VYCE(R)^GW^.

#### GoldenBraid based cloning

Vector used to generate CRISPR-Cas9 knockouts of *NbT5* was constructed using the GoldenBraid system, following the strategy described by Aliaga-Franco et al. (2019) (Aliaga-Franco *et al*., 2019). The sgRNA multiplex transcriptional unit was driven by the AtU6-26 promoter (GB1001). Transcriptional units for kanamycin resistance and humanized Cas9 (hCas9) were also included (EGM007). Additionally, a transcriptional unit for the DsRed fluorescent protein under the control of the 35S promoter (GB0361) was added for negative selection. All transcriptional units were assembled into the plant destination vector pDGB3α1 to generate the final plant expression vector.

### Transient expression in *N. benthamiana*

Transient expression in *N. benthamiana* leaves was performed using *Agrobacterium* strain GV3101::pMP90 carrying the appropriate constructs, along with the p19 silencing suppressor strain. Plants were grown for 21 days under long-day conditions (24 ⁰C, 65% humidity, 120 µmol photons m^−2^ s^−1^). *Agrobacterium* cultures were grown overnight in Luria-Bertani medium containing rifampicin (50 µg mL^-1^), gentamycin (25 µg mL^-1^) and construct-specific antibiotics. After growth, bacteria were harvested by centrifugation, pellets were resuspended in infiltration solution (10 mM MES pH 5.6, 10 mM MgCl_2_, and 1 mM acetosyringone) and incubated for 2 h in the dark at room temperature. For single construct experiments, *Agrobacterium* was infiltrated at OD_600_ of 0.6, the p19 strain at 0.3. For double infiltration experiments, *Agrobacterium* strains were infiltrated at OD_600_ of 0.4 for the constructs and 0.2 for the p19 strain. Leaves were infiltrated on the abaxial side and analyzed 2 days post infiltration.

### Confocal imaging

#### Subcellular localization/Bimolecular fluorescence complementation

Two days after infiltration, leaf disks from *N. benthamiana* were excised and visualized using an LSM 880 confocal microscope with a Plan-Apochromat 40x/1.2 W objective. GFP and mCherry were excited at 488 and 561 nm, respectively. Fluorophore detection involved a PMT, a GaAsP (used to improve signal detection), and an additional PMT for transmitted light. Cortical plane images are maximum Z-projections of several planes from the cell surface to a plane where cells are close but not in contact with neighboring cells (to facilitate individual cell identification). For BiFC experiments, reconstituted Venus fluorescence was excited at 514 nm and images were acquired with identical settings (laser intensity, detector gain, magnification) for all samples.

#### Chloroplast number analysis

Seedling cotyledons were analyzed using an LSM 880 confocal microscope with a Plan-Apochromat 40x/1.2 W objective. Chloroplasts were visualized through chlorophyll autofluorescence, and cells were observed using brightfield. Chlorophyll detection was enhanced with a GaAsP detector and a PMT was used for transmitted light. The number of chloroplasts per cell was counted in a single equatorial plane of each cell and normalized to the total cell area.

### Fixation and transmission electron microscopy imaging

Samples were processed for transmission electron microscopy as described by Sánchez-Sevilla et al., (2021). Briefly, cotyledons were fixed in 2.5% glutaraldehyde in 50 mM sodium cacodylate buffer (pH 7.4), post-fixed in 1% OsO₄, contrasted with 2% aqueous uranyl acetate, dehydrated through an ethanol series and embedded in London Resin White. Ultrathin sections (50–70 nm) were examined using a JEM-1400 transmission electron microscope.

### Image analysis

All images were analyzed using FIJI (Schindelin *et al*., 2012). For ER/chloroplast volume percentage, a macro was developed to quantify images (Supplementary File 3). Briefly, chloroplasts were visualized using chlorophyll autofluorescence. Chloroplast and ER were selected with auto local threshold (Bernsen method). Chloroplasts that were fragmented or not completed, as well as stomatal chloroplasts, were excluded before continuing to further analysis. A halo of 2 iterations through dilation was created surrounding each chloroplast to measure ER proximal to chloroplast. 3D object counter plugin was used to measure ER 3D volume in each halo. This was repeated for each chloroplast, and ER halo volume per chloroplast was calculated. 3D reconstruction of the images was built using MeshLab.

### Chlorophyll extraction and quantification

Total chlorophyll content in seedlings was extracted using methanol, following the method described by Warren (Warren, 2008). Briefly, approximately 50 mg of ground tissue per biological replicate was extracted with 1 mL of methanol, and the supernatant was recovered by centrifugation; pellets were re-extracted when necessary and supernatants pooled. Absorbance was measured at 652 and 665 nm in a microplate reader, and chlorophyll concentrations were calculated with the Ritchie formula (Ritchie, 2006) using a 1 cm corrected path length. Chlorophyll content in adult plants was measured using a Soil Plant Analysis Development (SPAD) chlorophyll meter. Each leaf was measured at four different locations, and the average value was calculated to estimate the overall chlorophyll content of the leaf.

### Total lipid extraction

Lipids were extracted from approximately 250 mg of *N. benthamiana* seedlings or 50 mg of seeds. The tissue was harvested, frozen in liquid nitrogen, and ground into a fine powder. Lipid extraction was performed using the Hara and Radin method (Hara and Radin, 1978), using glassware throughout. Briefly, 8 volumes (v/w) of isopropanol were added to the samples and heated at 80 °C for 5 min. After cooling, 12 volumes (v/w) of hexane were added and mixed. Subsequently, 10 volumes (v/w) of 6.7% sodium sulfate were added. The mixture was thoroughly mixed and centrifuged at 21,000g for 5 min to separate the lipid extract from the tissue. The upper phase was transferred to a clean tube, and the pellet was re-extracted with 15 volumes (v/w) of hexane:isopropanol (7:2). The lipid extracts (upper phases) were combined. The solvents were evaporated under nitrogen gas until approximately 1 mL of solvent remained.

### Statistical analysis

Data were analyzed using GraphPad Prism 8.02 (GraphPad Software). The selection of the statistical test was based on the type of data, the number of groups compared, the results of the normality test, and the differences in standard deviations between samples. Normality was assessed using the Shapiro-Wilk test. The statistical test applied to each sample set is specified in the corresponding figure legend.

## SUPPLEMENTARY DATA DESCRIPTION

**Supplementary file 1**. Sequences used in Fig. 1(A-C).

**Supplementary file 2.** Sequences used in Fig S1D.

**Supplementary file 3.** ImageJ macro used to quantify ER surrounded each chloroplast.

## ETHICS APPROVAL AND CONSENT TO PARTICIPATE

Not applicable

## CONSENT FOR PUBLICATION

Not applicable

## AVAILABILITY OF DATA AND MATERIALST

All data supporting the findings of this study are available within the article and their supplementary materials.

## AUTHORS’ CONTRIBUTIONS

**Carolina Huercano**: Conceptualization, Formal Analysis, Investigation, Visualization, Writing-original draft. **Oliver Cuevas:** Investigation, Visualization, Writing-review & editing. **Paula Velasco-Palomo:** Investigation, Visualization, Writing-review & editing. **Miriam Moya-Barrientos:** Investigation, Visualization, Writing-review & editing. **Francisco Percio:** Investigation, Visualization, Writing-review & editing. **Joaquin J. Salas-Liñan:** Investigation, Writing-review & editing. **Victoria Sanchez-Vera:** Conceptualization, Writing-review & editing, **Noemi Ruiz-Lopez:** Conceptualization, Funding acquisition, Project administration, Resources, Supervision, Writing-original draft.

## ACKNOWLEDGMENTS

We thank Alicia Esteban at the microscopy facilities at the IHSM for assistance with confocal microscopy and Manuela Vega Sánchez from Servicio Central de Informatica from Universidad de Málaga for her assistance with macro development and image analysis.

## ABBREVIATIONS

35S: cauliflower mosaic virus 35S promoter
At: Arabidopsis thaliana
a.u.: arbitrary units
AtT5.1: AtNTMC2T5.1
AtT5.2: AtNTMC2T5.2
BAG7: BCL-2-associated athanogene 7
BiFC: bimolecular fluorescence complementation
C2: Ca²⁺-binding domain
CaLB: calcium-dependent lipid-binding
CLANS: Cluster Analysis of Sequences
DAG: diacylglycerol
DAS: days after sowing
DGDG: digalactosyldiacylglycerol
ER: endoplasmic reticulum
ERMES: ER–mitochondria encounter structure
E-Syt: extended synaptotagmin
FFA: free fatty acids
FW: fresh weight
GFP: green fluorescent protein
hCas9: human codon-optimized
Cas9 HR: hydrophobic region
Hs: Homo sapiens
kanR: kanamycin resistance gene
LB: left border
MGDG: monogalactosyldiacylglycerol
MS: Murashige and Skoog
Nb: Nicotiana benthamiana
NbT5: NbNTMC2T5
NTMC2T: N-terminal-transmembrane-C2 domain type
OMP24: outer membrane protein of 24 kDa
ORP2A: oxysterol-binding protein-related protein 2A
PC: phosphatidylcholine
PDZD8: PDZ domain-containing protein 8
PE: phosphatidylethanolamine
PG: phosphatidylglycerol
PI: phosphatidylinositol
Sc: Saccharomyces cerevisiae
SFH5/SFH7: Sec14 homolog 5/7
SMP: synaptotagmin-like mitochondrial-lipid-binding protein
SPAD: soil plant analysis development
SQDG: sulfoquinovosyldiacylglycerol
SYT: synaptotagmin
TAG: triacylglycerol
Tcb: tricalbin
TEM: transmission electron microscopy
TEX2: testis-expressed protein 2
TGD: trigalactosyldiacylglycerol
TM: transmembrane region
TMEM24: transmembrane protein 24
TULIP: tubular lipid-binding
UBQ10: UBIQUITIN10
VAP27: vesicle-associated membrane protein-associated protein 27
VYCE: C-terminal fragment of Venus
VYNE: N-terminal fragment of Venus
WT: wild type

## SUPPLEMENTARY FIGURES

**Figure S1.**
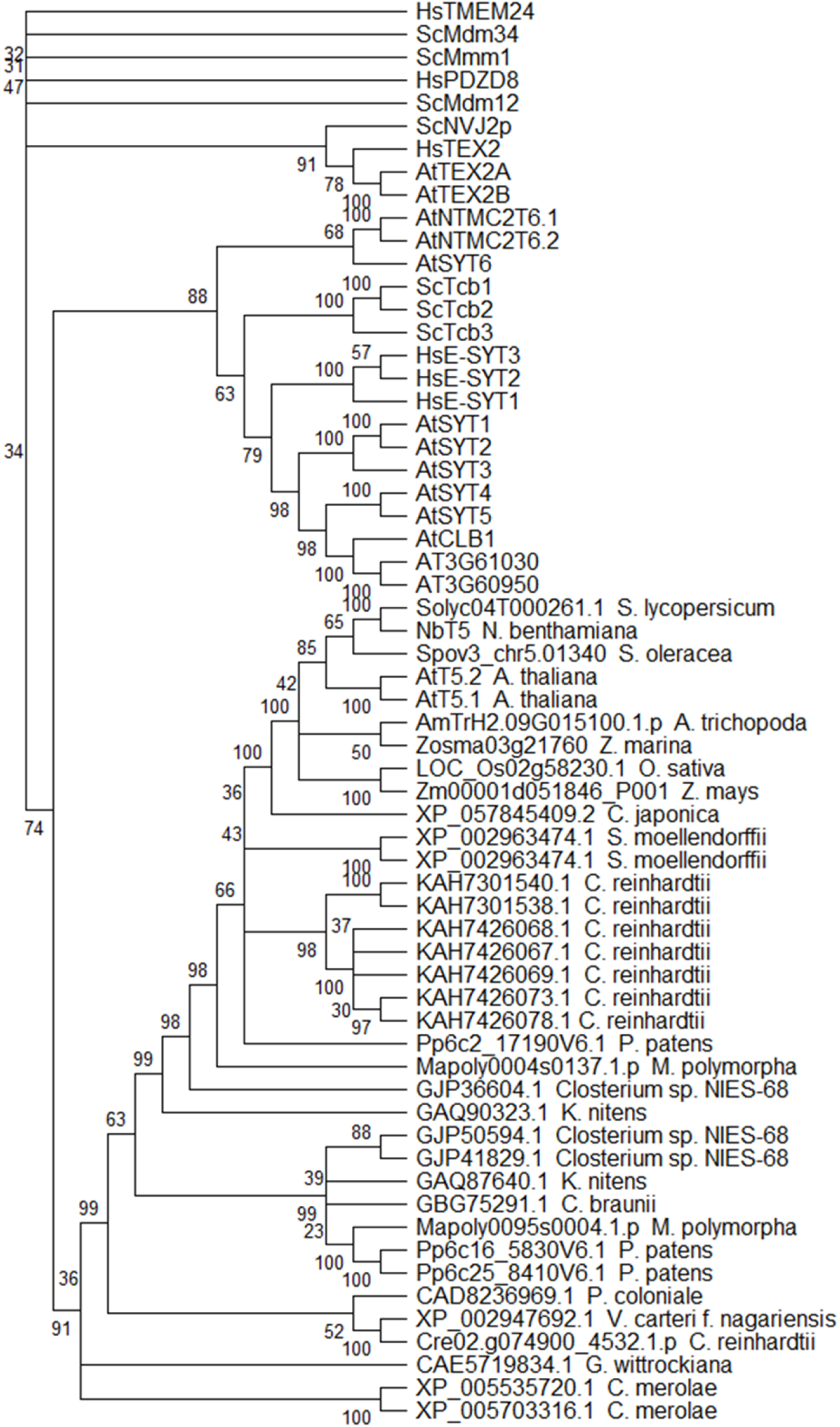
Evolutionary analysis of the NTMC2T5 proteins across the Archaeplastida lineage. Bootstrap consensus tree of NTMC2T5 family proteins identified from red algae to eudicot species, together with previously characterized SMP family proteins from *Homo sapiens*, *Saccharomyces cerevisiae*, and *Arabidopsis thaliana*. Candidate homologs were identified by PSI-BLAST and filtered to retain only proteins with the conserved TM– SMP–C2–HR domain architecture of the *A. thaliana* and *Nicotiana benthamiana* NTMC2T5 proteins (Fig. 2A). Sequences were aligned with MAFFT v7 (Katoh *et al*., 2019) using default settings, and the tree was reconstructed by maximum likelihood in MEGA12 (Kumar *et al*., 2024) under the LG+G+I+F model, as described in Methods. Bootstrap values from 500 replicates are indicated next to the corresponding branches; branches reproduced in less than 50% of replicates were collapsed. The protein sequences used are provided in Supplemental File S2. *Abbreviations:* AtT5.1, AtNTMC2T5.1; AtT5.2, AtNTMC2T5.2; NbT5, NbNTMC2T5.

**Figure S2.**
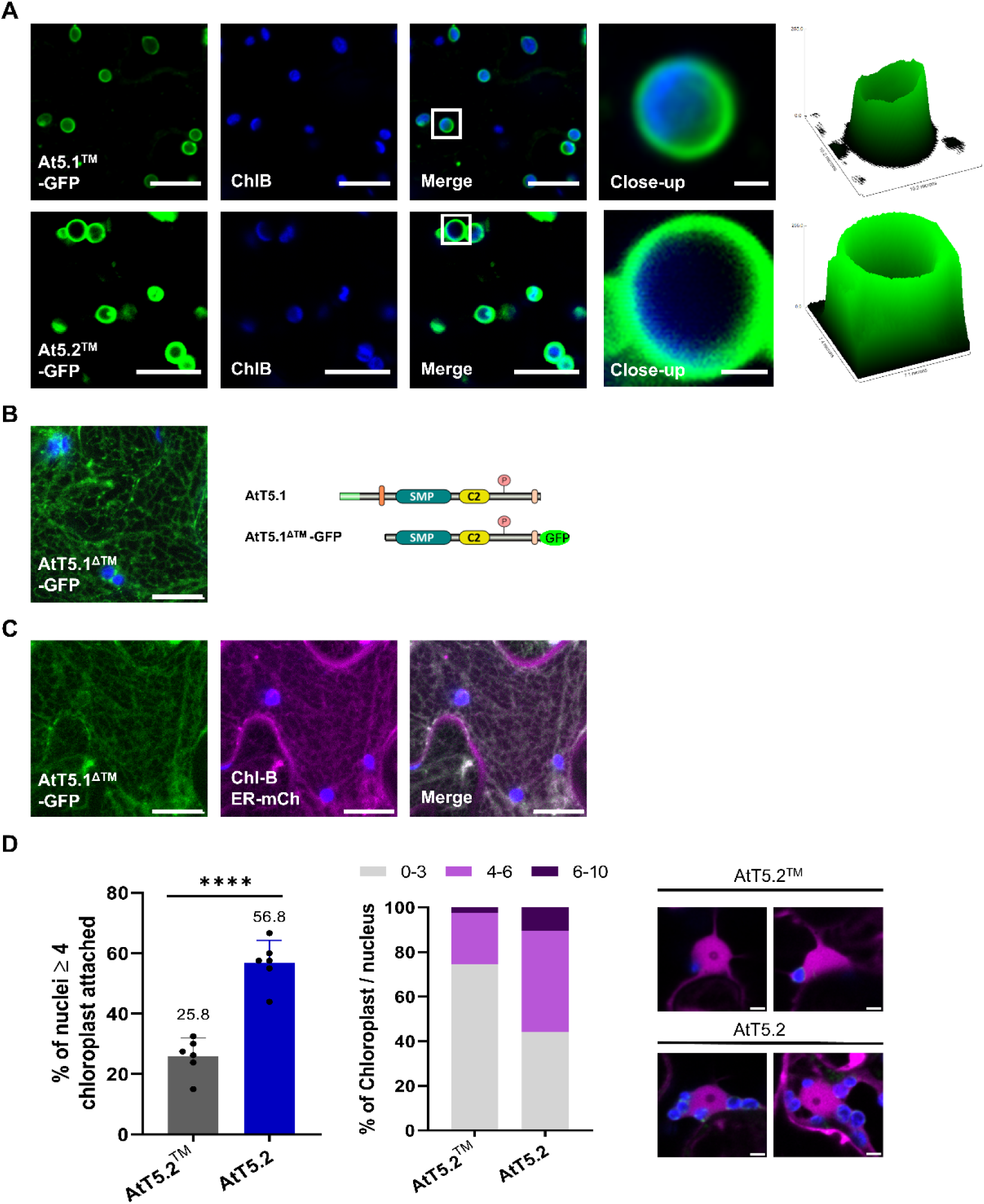
The transmembrane region determines chloroplast targeting of Arabidopsis NTMC2T5 proteins, and NTMC2T5.2 induces chloroplast clustering in *Nicotiana benthamiana*. **(A)** Maximum Z-projections of confocal images of *N. benthamiana* epidermal cells transiently expressing AtT5.1^TM^-GFP (M1-G160) or AtT5.2^TM^-GFP (M1-G179). Both proteins displayed a uniform distribution around chloroplasts. Boxed regions are magnified in the insets. Surface plots of the fluorescence intensity of the insets (right panels) illustrate the spatial distribution of the GFP signal. **(B)** Maximum Z-projections of confocal images of *N. benthamiana* epidermal cells transiently expressing AtT5.1^ΔTM^-GFP (E183-P675). The GFP signal displayed a reticulate pattern. **(C)** Maximum Z-projections of confocal images of *N. benthamiana* epidermal cells co-expressing AtT5.1^ΔTM^-GFP and ER marker-mCh. The GFP signal partially overlapped with the ER network. **(A–C)** Chloroplasts were visualized by chlorophyll autofluorescence (blue), GFP fluorescence is shown in green and mCherry fluorescence in magenta. Individual channels and the merged image are shown. Scale bars: 20 µm; inset scale bars: 2 µm. **(D)** Quantification of chloroplast clustering around the nucleus in epidermal cells of *N. benthamiana* leaves transiently expressing AtT5.2^TM^-GFP (M1-G179) or AtT5.2-GFP. Free mCherry was co-expressed in all cases to visualize nuclei. From left to right: percentage of nuclei with ≥ 4 attached chloroplasts, number of chloroplasts per nucleus, and representative single-plane confocal images containing the equatorial section of the nucleus for each condition. Merged images show chlorophyll autofluorescence (blue), GFP (green) and free mCherry (magenta). Bars represent mean ± SD. Six plants per condition were analyzed, with approximately 20 nuclei per plant. Data were analyzed using an unpaired two-tailed Student’s t-test. Asterisks indicate significant differences (ns, not significant; ****, *P* < 0.0001). Scale bar: 5 µm. *Abbreviations: AtT5.1, AtNTMC2T5.1; AtT5.2, AtNTMC2T5.2; HR, hydrophobic region; TM, transmembrane region*.

**Figure S3.**
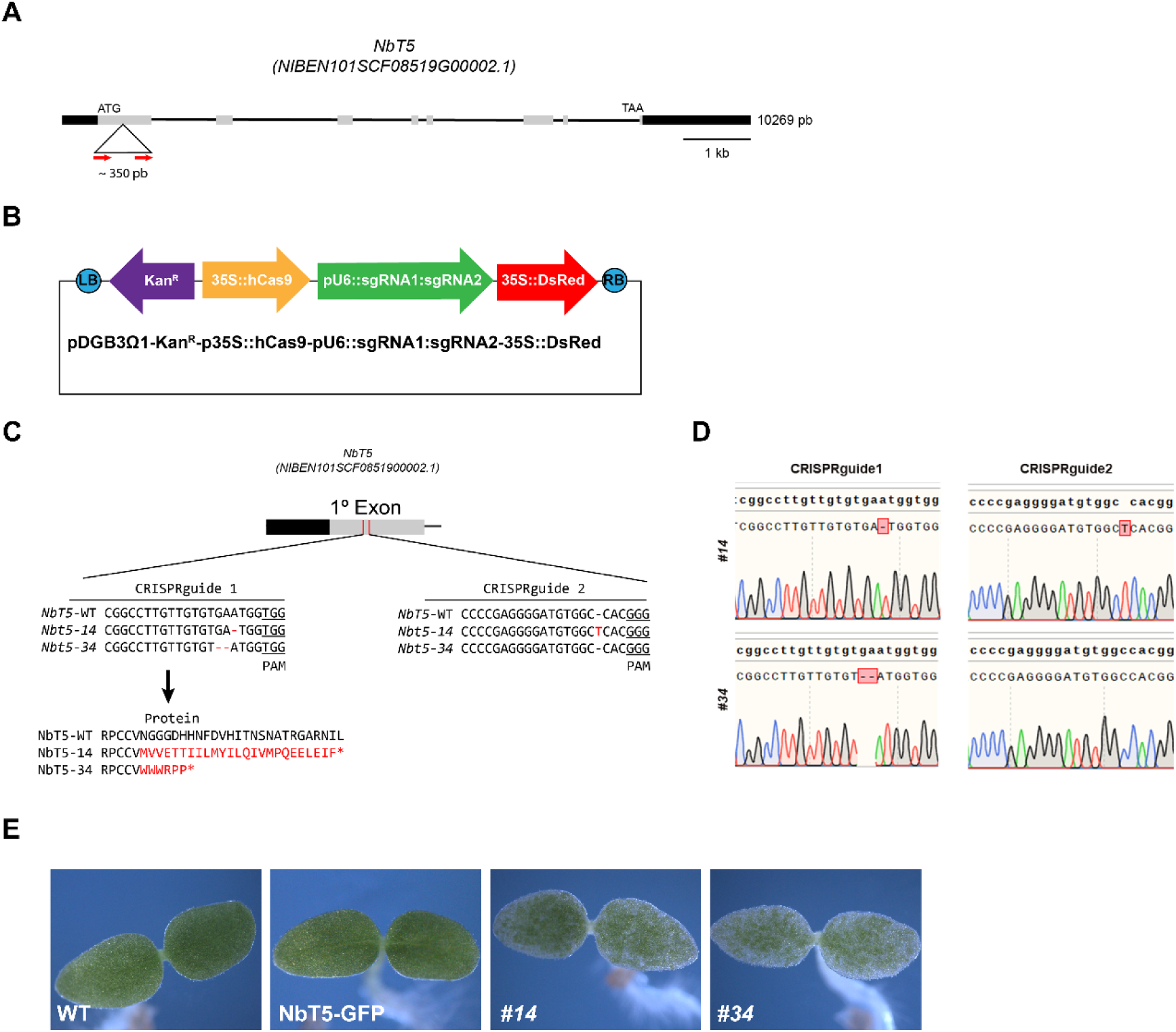
Generation and validation of CRISPR/Cas9-edited and complemented Nicotiana NTMC2T5 lines. **(A)** Schematic representation of the *NbT5* gene structure. Boxes represent coding exons and lines represent introns. Red arrows indicate the positions of the two sgRNAs used for CRISPR/Cas9-mediated mutagenesis. **(B)** Diagram of the GoldenBraid CRISPR/Cas9 construct used for plant transformation. **(C)** Sequence alignment showing mutations detected in two homozygous *nbt*5 lines (*#14* and *#34*); the mutation carried by each line is indicated in the figure. Mismatches are indicated by red dashes. The predicted protein products of both mutants are also shown. **(D)** Sequencing chromatograms of the *nbt5* lines *#14* and *#34*. Red boxes highlight mismatches between WT and mutant sequences. **(E)** Stereomicroscope images of cotyledons from WT, *nbt5* mutants *#14* and *#34*, and NbT5-GFP lines at 5 days after sowing. *Abbreviations:* 35S, cauliflower mosaic virus (CaMV) 35S promoter; hCas9, human codon-optimized Cas9; kan^R^, kanamycin resistance gene; LB, left border; NbT5, NbNTMC2T5; pU6, U6 promoter; RB, right border; sgRNA, single-guide RNA.

**Figure S4.**
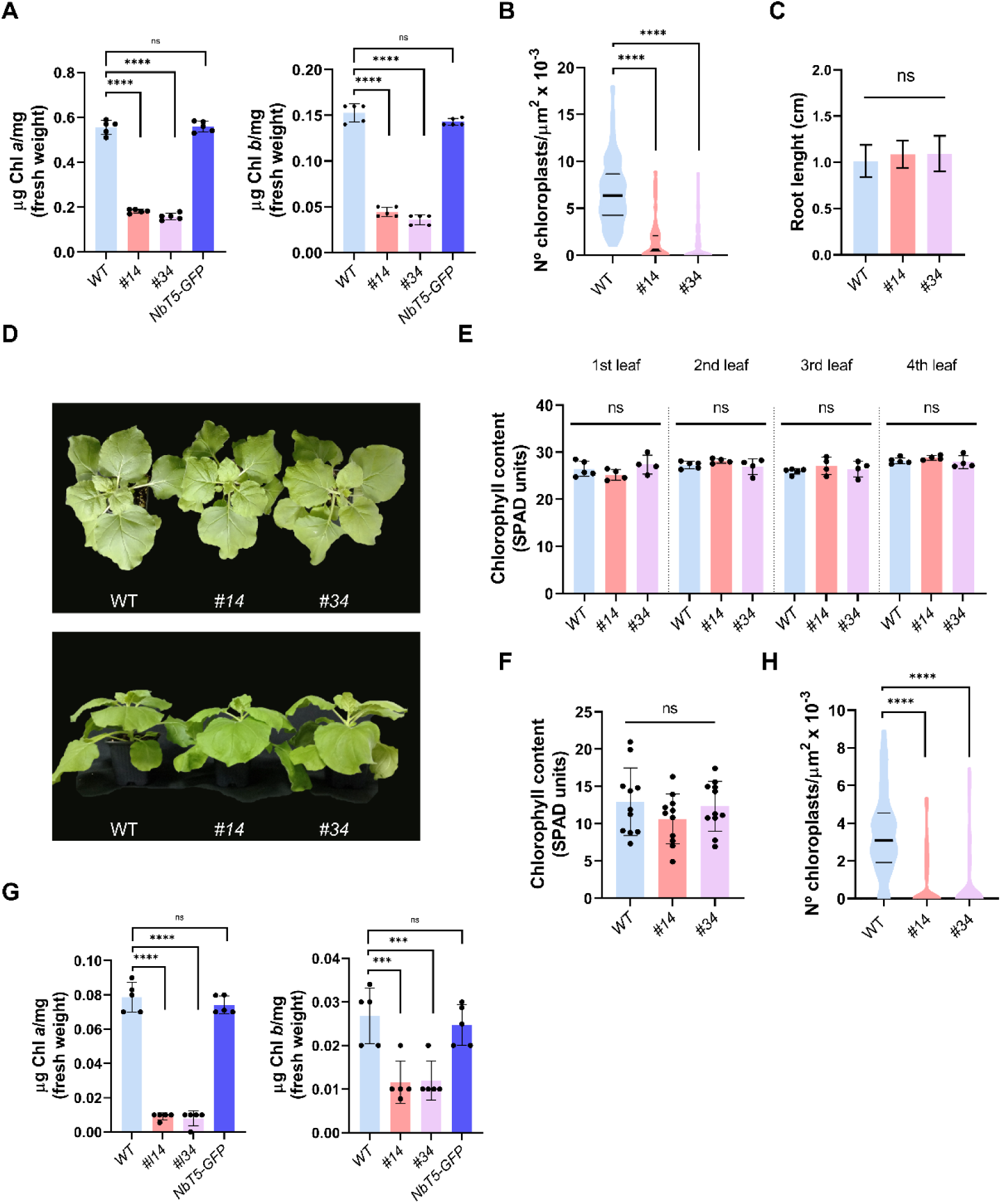
Developmental characterization of *Nicotiana benthamiana ntmc2t5* plants reveals chloroplast-associated phenotypes during early seedling development but not in adult stages. **(A)** Chlorophyll *a* and chlorophyll *b* contents in 7-day-old seedlings grown under long-day conditions. **(B)** Chloroplast density in cotyledon cells of 7-day-old seedlings grown under long-day conditions on MS medium supplemented with 1.5% sucrose. Chloroplast density was calculated for individual cells by dividing the total number of chloroplasts within each cell by its measured cell area (chloroplasts µm⁻²). **(C)** Root length of WT and *nbt5* lines *#14* and *#34* measured 7 days after sowing. **(D)** General shoot appearance of 4-week-old WT and *nbt5* mutant plants. **(E)** Chlorophyll content (SPAD units) of the first, second, third and fourth leaves after the cotyledons in 4-week-old plants. **(F)** Chlorophyll content (SPAD units) of senescent leaves in 7- to 8-week-old WT and *Nbt5* plants. The first leaf after the cotyledons was measured. **(G, H)** Seedlings were grown in complete darkness for 7 days and then transferred to long-day conditions for 24 h before analysis. **(G)** Chlorophyll *a* and chlorophyll *b* contents in de-etiolated seedlings. **(H)** Chloroplast density in cotyledon cells of de-etiolated seedlings grown on MS medium supplemented with 1.5% sucrose, quantified as in (B). In (A), (C), (E), (F) and (G), bars represent the mean ± SD, and data were analyzed by one-way ANOVA followed by Dunnett’s multiple comparison test relative to WT. Data in (B) and (H) are shown as violin plots and were analyzed using the Kruskal-Wallis test followed by Dunn’s multiple comparison test. Asterisks indicate statistical significance (ns, not significant; ***, *P* ≤ 0.001; ****, *P* ≤ 0.0001). *Abbreviations:* Chl, chlorophyll; *NbT5, NbNTMC2T5*.

**Figure S5.**
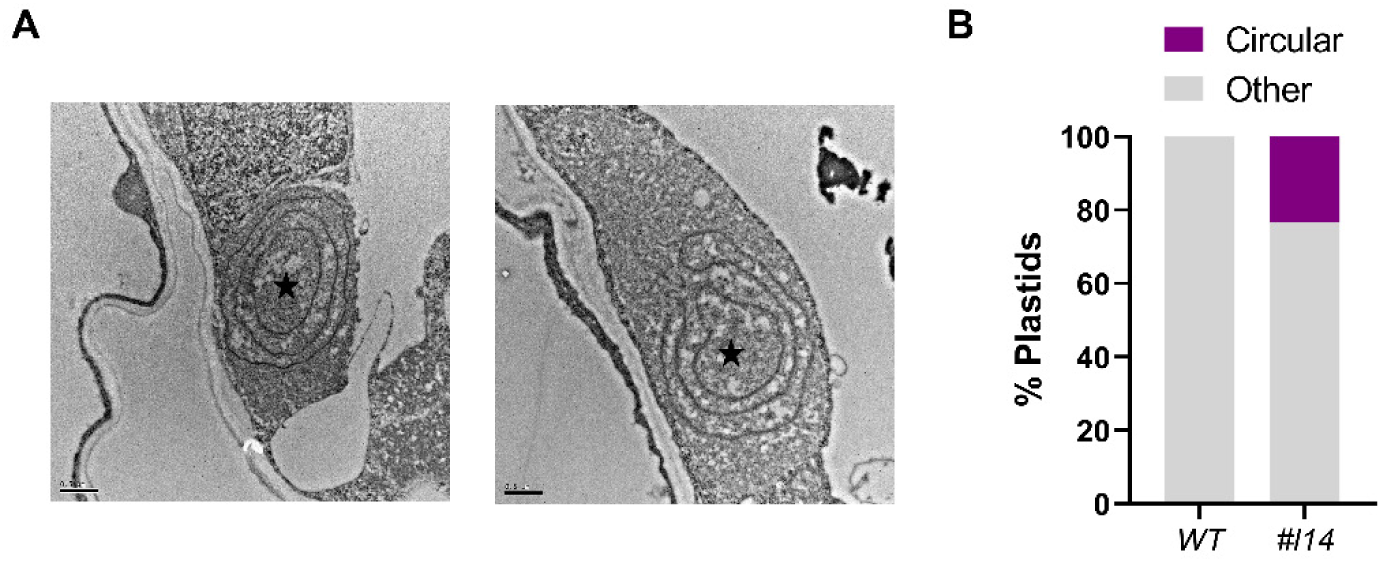
Circular prothylakoid-like structures in plastids of *ntmc2t5* cotyledons. **(A)** Transmission electron micrographs of cotyledons from WT and *nbt5 #14* seedlings grown for 7 days in darkness followed by 1 day in light. Black stars indicate circular lamellar prothylakoids within proplastid-like structures. Scale bar: 500 nm. **(B)** Percentage of plastids containing circular prothylakoid-like structures (n = 33 plastids per genotype). *Abbreviations: nbt5, nbntmc2t5*.

**Figure S6.**
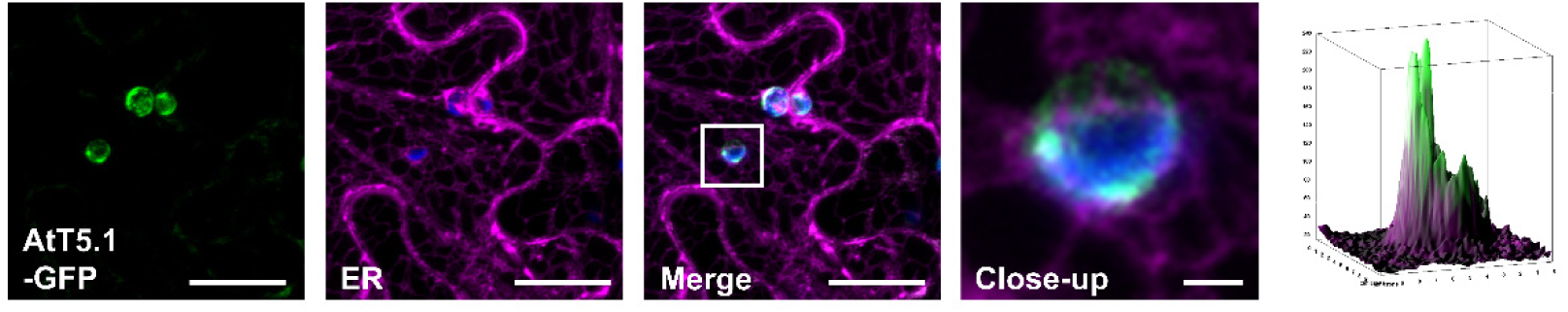
Arabidopsis NTMC2T5.1-GFP shows partial co-localization with the ER network in *N. benthamiana*. Maximum-intensity Z-projections of confocal images of *N. benthamiana* leaves transiently co-expressing AtT5.1-GFP and an ER marker (ER-mCh). The boxed region is magnified in the inset. Surface plots of the fluorescence intensity (right panel) illustrate the spatial distribution of the signals within the inset. Chloroplasts were visualized by chlorophyll autofluorescence (blue), and GFP and ER-mCh fluorescence are shown in green and magenta, respectively. Individual channels and the merged image are shown. Scale bar: 20 µm; inset scale bar: 2 µm. *Abbreviations:* AtT5.1, AtNTMC2T5.1.

**Figure S7.**
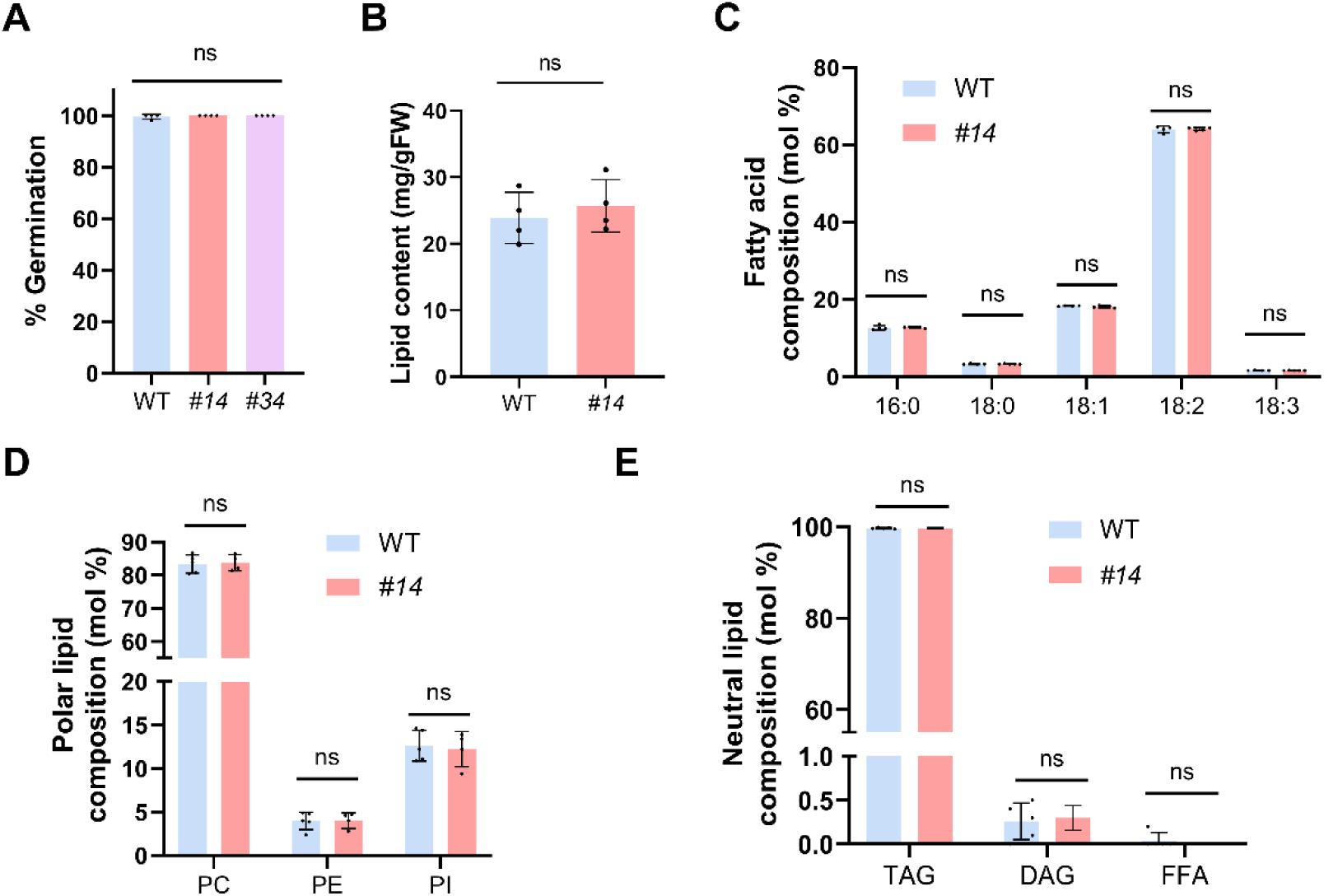
**Ntmc2t5 mutants show no significant changes in seed germination or lipid composition. (A)** Germination rates of WT and *nbt5* lines *#14* and *#34*. Seeds were plated on ½ MS medium and scored after 3 days at 25 °C. Germination percentages were calculated based on radicle emergence, using four sets of 200 seeds each. **(B-E)** Lipid analysis of seeds from WT and *nbt5* line *#14*. **(B)** Total seed lipid content (mg g⁻¹ fresh weight). **(C)** Seed fatty acid distribution (mol %). **(D)** Polar lipid composition (mol %). **(E)** Neutral lipid composition (mol %). **(A–E)** Bars represent the mean ± SD. Data in (A) were analyzed using the Kruskal-Wallis test followed by Dunn’s multiple comparison test. Data in (B) were analyzed by unpaired two-tailed Student’s *t*-test, and data in (C– E) by two-way ANOVA followed by Sidak’s multiple comparison test. No significant differences were detected in any of the comparisons (ns, not significant). *Abbreviations*: DAG, diacylglycerol; FFA, free fatty acids; FW, fresh weight; *nbt5, nbntmc2t5*; PC, phosphatidylcholine; PE, phosphatidylethanolamine; PI, phosphatidylinositol; TAG, triacylglycerol.

**Table S1.**
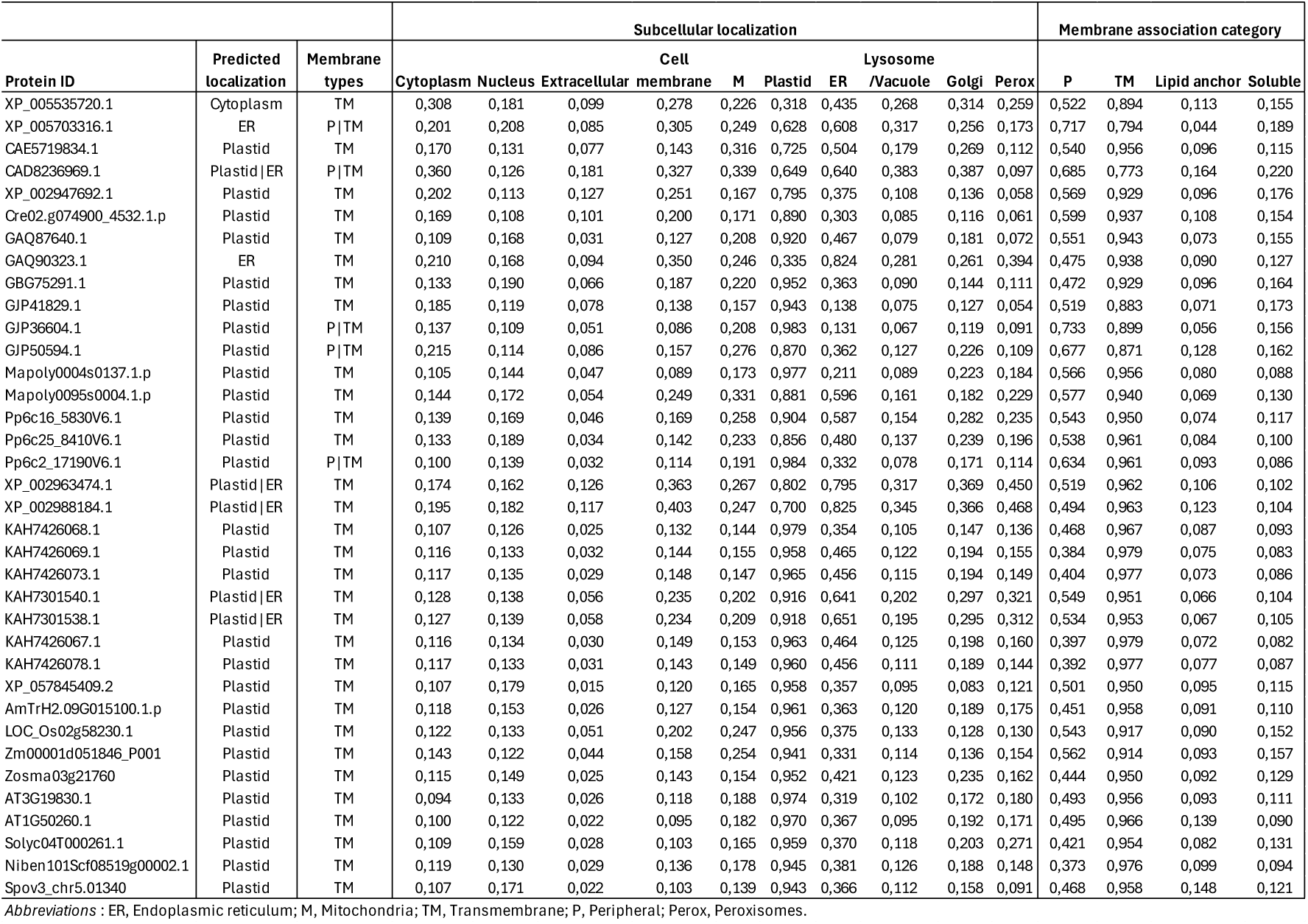
DeepLoc 2.1 prediction of the subcellular localization of the putative NTMC2T5 proteins identified across the Archaeplastida lineage.

**Table S2.**
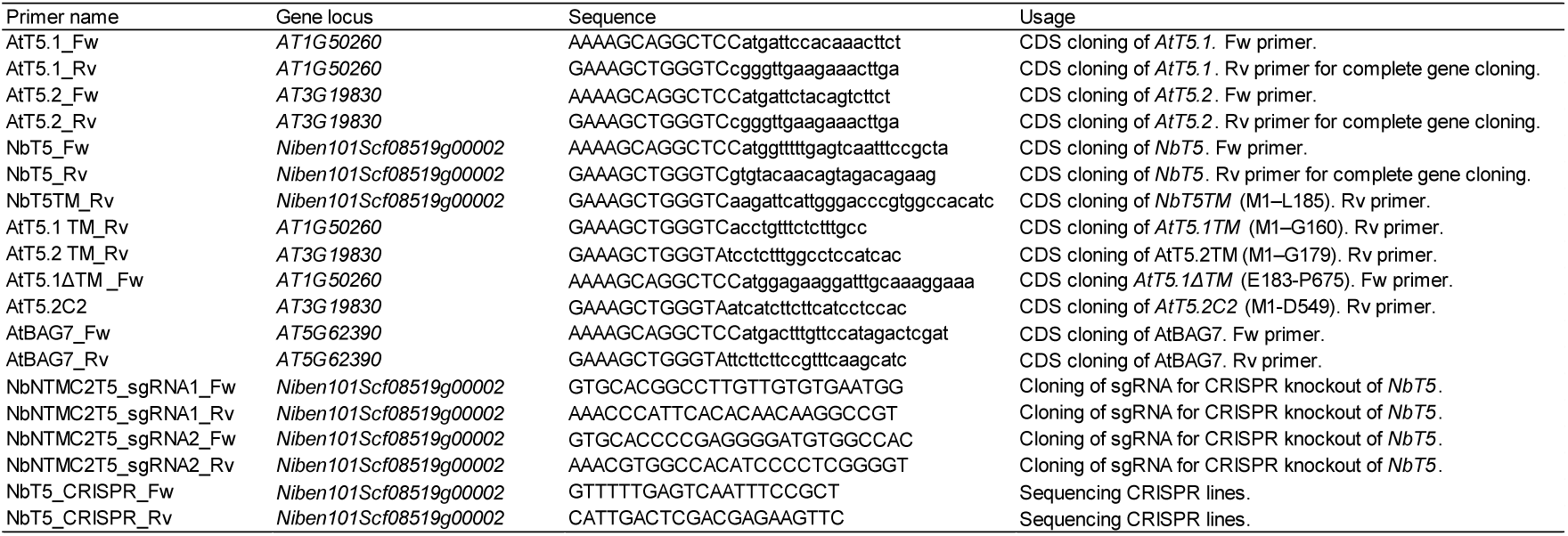
List of primers used in this study. Uppercase letters correspond to *att*B sites for Gateway cloning.

**Table S3.**
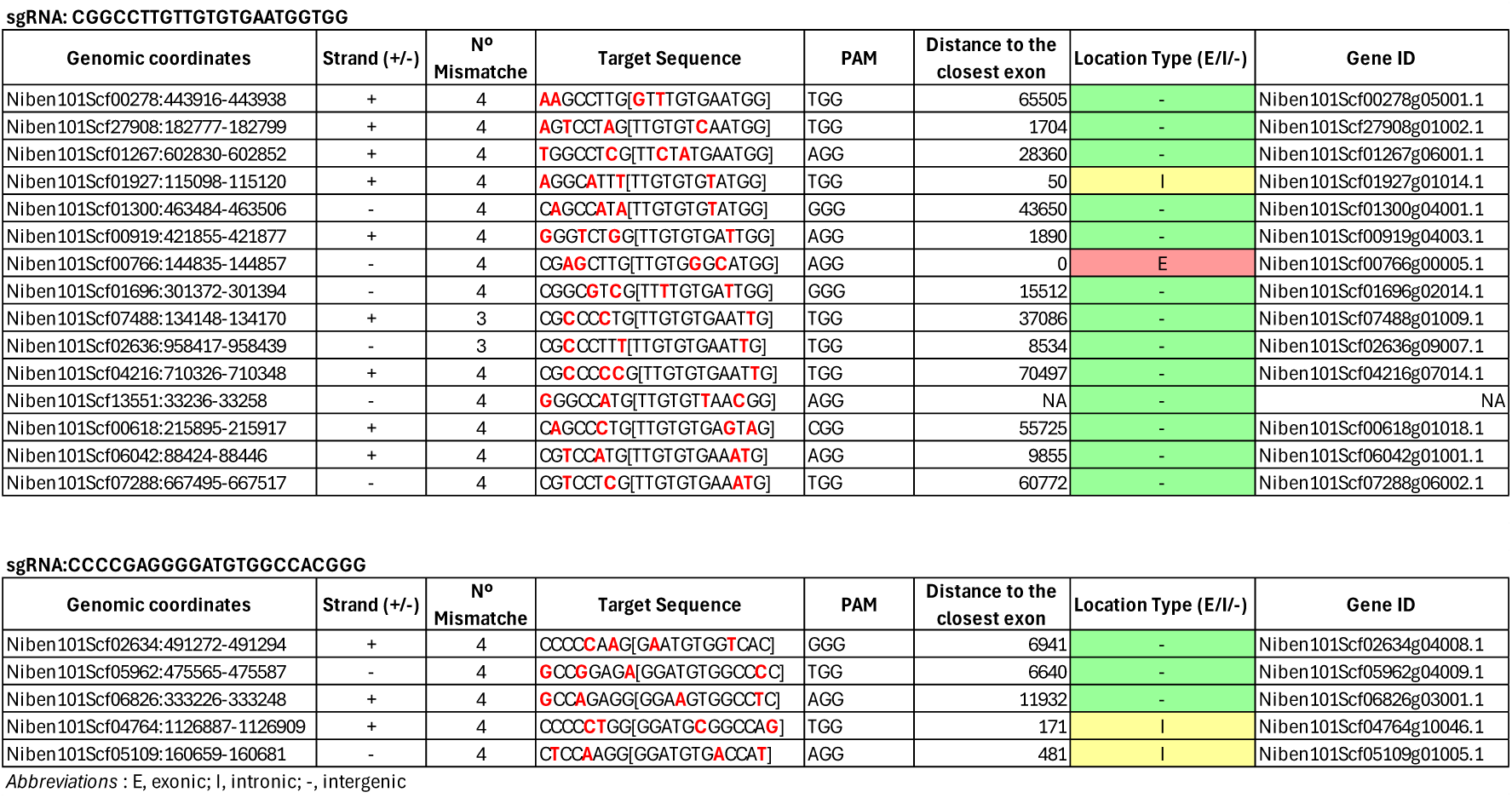
Computational prediction of potential CRISPR/Cas9 off-target sites using CCTop.

